# A Highly Selective Chemical Probe for Activin Receptor-like Kinases ALK4 and ALK5

**DOI:** 10.1101/2020.01.23.916502

**Authors:** Thomas Hanke, Jong Fu Wong, Benedict-Tilmann Berger, Ismahan Abdi, Lena Marie Berger, Roberta Tesch, Claudia Tredup, Alex N. Bullock, Susanne Müller, Stefan Knapp

## Abstract

The transforming growth factor beta-receptor I/activin receptor-like kinase 5 (TGFBR1/ALK5) and its close homologue ALK4 are receptor protein kinases associated with the development of diverse diseases, including cancer, fibrosis, heart diseases and dysfunctional immune response. Therefore, ALK4/5 are among the most studied kinases and several inhibitors have been developed. However, current commercially available inhibitors either lack selectivity or have not been comprehensively characterized, limiting their value for studying ALK4/5 function in cellular systems. To this end, we report the characterization of the 2-oxo-imidazopyridine, TP-008, a potent chemical probe with dual activity for ALK4 and ALK5 as well as the development of a matching negative control compound. TP-008 has excellent cellular potency and strongly abrogates phosphorylation of the substrate SMAD2 (mothers against decapentaplegic homolog 2). Thus, this chemical probe offers an excellent tool for mechanistic studies on the ALK4/5 signaling pathway and the contribution of these targets to disease.

## Introduction

The transforming growth factor-*β* (TGF-*β*) superfamily of signaling ligands is a large family, which comprises in mammals at least 30 structurally related cytokines including the TGF-*β*s, bone morphogenetic proteins (BMPs), activins, growth and differentiation factors (GDFs), the proteins nodal and the anti-Muellerian hormone (AMH).*(1)* Because of its involvement in many cellular processes such as proliferation, cell growth, adhesion, migration, differentiation and other regulatory functions, the TGF-*β* signaling pathway has been widely studied. *(1,2)*

TGF-β signaling is initiated by the binding of a dimeric TGF-β ligand which assembles a heterotetrameric transmembrane serine-threonine kinase receptor complex, consisting of two TGF-β type II receptors (TGFBR2) and two type I receptors (TGFBR1/ALK5, also known as TβRI) *(3)*. Once this heterotetrameric complex is formed, ALK5 is activated and phosphorylates the intracellular regulatory SMADs (R-SMADs). R-SMADs in turn bind to the common SMAD (Co-SMAD) SMAD4, inducing their translocation into the nucleus, where they promote the expression of specific target genes (Figure 1A). Dysregulated expression of genes linked to this pathway has been associated with the development of cancer, fibrosis, immunity or heart diseases *(4–7)*. Consequently, inhibition of TGF-β receptor superfamily has gained increasing attention over the last two decades, but so far no drug inhibiting TGF-β family receptors has been approved. Five type II and seven type I receptors, also termed activin-receptor like kinases (ALK1-7), have been identified in mammals, which have a similar sequence and domain structure composed of a ligand-binding extracellular domain, a transmembrane domain, and a cytoplasmic kinase domain *(8)*. They belong to the serine/threonine kinase receptor (STKR) subfamily, which is part of the tyrosine kinase-like (TKL) branch of kinases. ALK5/TGFBR1 and ALK4/ACVR1B are both closely related and phosphorylate SMAD2 and SMAD3, whereas ALK1/2/3 and ALK6 phosphorylate SMAD1/5/8.

**Figure 1.**
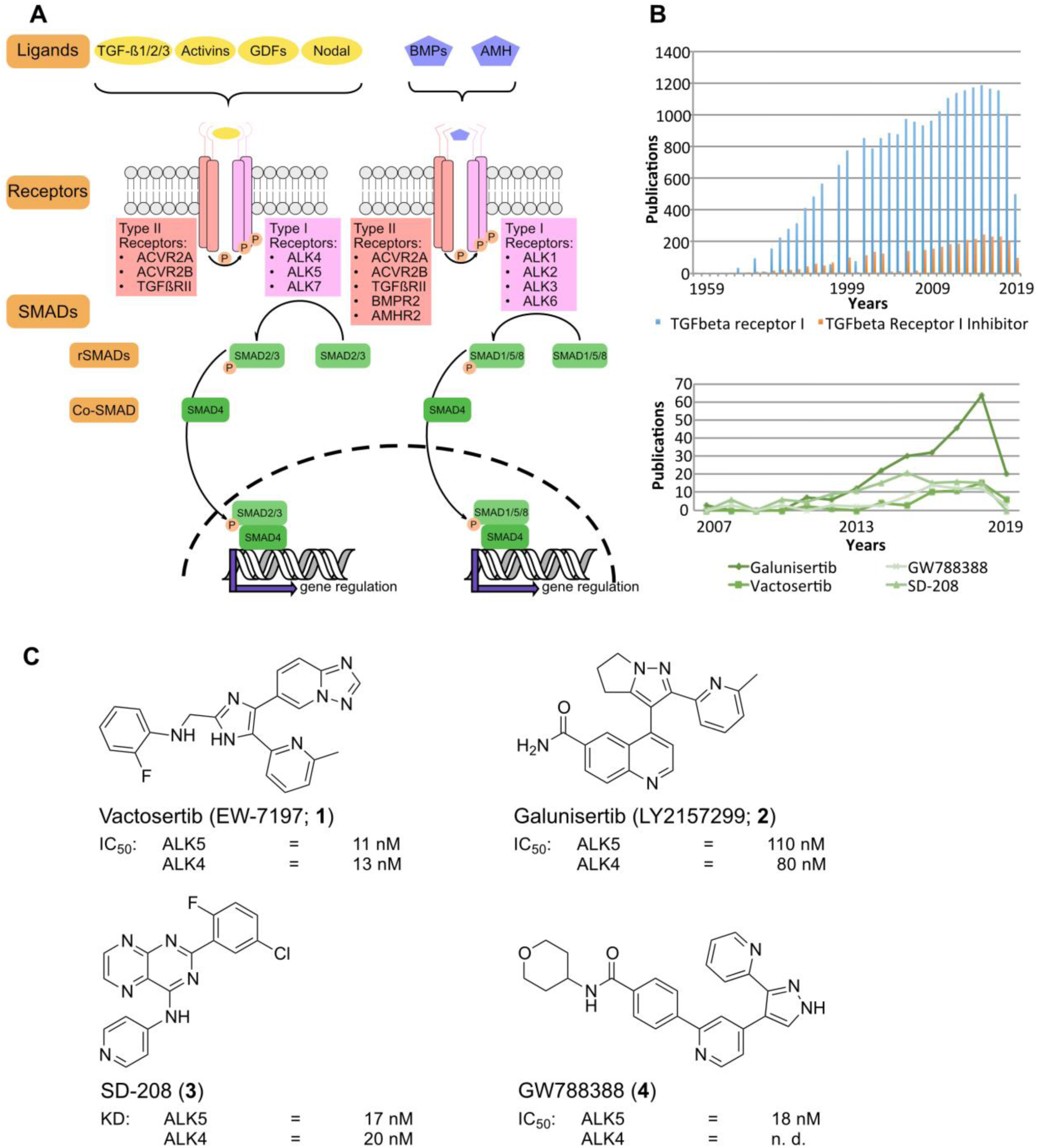
(A) Signaling pathways of the TGF-β/SMAD family. (B) Publications on ALK5 and ALK5 inhibitors over the last decades. Upper panel shows the overall publications with search term “TGFbeta receptor I” or “TGFbeta Receptor I + inhibitor” in Pubmed. Lower panel shows the number of publication using one of these inhibitors (**1–4**). (C) Most frequently used commercially available ALK5 inhibitors and their *IC*_50_ values on ALK4/5.

Several ALK4/5 inhibitors such as the clinical inhibitors Vactosertib (EW-7197; **1**) and Galunisertib (**2**), or the preclinical inhibitors, SD-208 (**3**) or GW-788388 (**4**) have been developed (Figure. 1C). Galunisertib is the most advanced ALK5 inhibitor which has now progressed to phase II clinical studies for treatment of different cancers *(9,10)*. However, current inhibitors have insufficient selectivity for mechanistic studies, inhibiting five to ten additional kinases with significant activity. For instance, Vactosertib (**1**) also inhibits RIPK2 and VEGFR2 with *IC*_50_ values below 100 nM and Galunisertib (**2**) inhibits besides ALK4/5 RIPK2, TGFBR2, MINK, CSNK1A1, MAP4K4, GAK, CSNK1E1 and BMPR1B with *IC*_50_ values ranging from 190 nM to 470 nM *(11,12)*. Recently Boehringer Ingelheim launched on the opnMe platform the compound BI-4659 to use as an ALK5 inhibitor. However also for this compound several off-targets have been described *(13,14)*. Thus, there remains a pressing need to develop a more selective, potent and cell active ALK4/5 inhibitor. Analysis of the literature revealed that more than 50% of published studies in 2018, used ALK4/5 inhibitors, involved one of the compounds **1–4**, whose selectivity profiles either reveal significant off targets (**1**–**3**) *(11,12)* or lack comprehensive characterization (**4**) (Figure 1C). Therefore, a chemical probe for this key target and important signaling molecule is of high interest for the research community. We and others have defined stringent criteria for chemical probes. A suitable tool compound for protein kinase research should display a biochemical potency below 100 nM at *K*_*M*_ of ATP, a cellular potency below 1 µM and a selectivity profile with no other off-targets within 30-fold apart potency from closely related paralogues *(15,16)*. Ideally, the chemical probe is also accompanied by a negative control compound, which is structurally related, but inactive against the target of interest. To increase the availability of chemical probes, the Structural Genomics Consortium (SGC) and its partners established the Donated Chemical Probe program through which high-quality compounds from the pharmaceutical industry or academic research laboratories are donated to the scientific community, sometimes following additional characterization and development of a suitable control compound *(17)*. Recently, a series of inhibitors including compound TP-008 (**5**) was reported by Takeda as a potent ALK5 inhibitor with an *IC*_50_ value of 25 nM and a promising pharmacokinetic profile *(18)*. TP-008 (**5**) shows no CYP inhibition (CYP1A2, CYP3A4), a low clearance and a low hERG liability (21% inhibition @ 10 µM). Initial selectivity profiling at 1 µM in a panel of about 50 kinases showed favorable selectivity and revealed no significant off-targets *(18)*. Here, we report the comprehensive characterization of TP-008 as a chemical probe for ALK4/5 and the development of a suitable negative control.

## Results and Discussion

In order to assess the selectivity of compound **5** (TP-008) in a wider kinase assay panel, TP-008 was screened in the *scan*Max_SM_ Kinase Assay Panel by Eurofins (DiscoverX) against 469 kinases *(19,20)*. No off-target activity was observed for TP-008 (**5**) at a screening concentration of 1 µM, while ALK4 and ALK5 exhibited 3.3% and 3.6% residual binding of the tracer molecule in this displacement assay format, confirming that TP-008 is a highly selective dual ALK4/5 inhibitor (Figure 3A, Table S5). Next, we tested the activity of TP-008 against both kinases in an enzyme kinetic kinase assay to determine the *in vitro* potency on ALK4 and ALK5 *(21)*. In comparison, Vactosertib (**1**) and GW-788388 (**4**) were also investigated. All assays were performed at an ATP concentration of 1 µM. Interestingly, TP-008 showed similar activity on both kinases with comparable *IC*_50_ values of 113 nM against ALK4 and 343 nM against ALK5, respectively, (Figure 2A) in good agreement with the results from the *scan*Max_SM_ Kinase Assay (Figure 3). In comparison, GW-788388 and Vactosertib showed *IC*_50_ values against ALK4/5 in the same assay of 155/222 nM and 8.3/11.8 nM, respectively (Table S1, Figure S1 and S2). Sequence alignment of ALK4 and ALK5 demonstrated a sequence identity of ∼70% and an identity of greater than 90% in the ATP-binding pocket (Tables S2-S4). The glycine rich loop, αC, β4, β5, hinge-region and αD especially were noted to share identical amino acid residues, indicating that selectivity between these two kinases may be difficult to achieve, if not impossible for an ATP competitive kinase inhibitor (type I or II). In contrast, one of the major differences of the type I receptors (ALK1–3 and ALK6) over ALK4/5 is the somewhat bulkier gatekeeper (threonine vs. serine), which has been explored to achieve selectivity between ALK subfamilies. However, the closest member to ALK4/5, within the type I receptors, is ALK7, which also harbors a serine gatekeeper and differs just in one position in the hinge region (D281 in ALK5 vs. E271 in ALK7), the αD-Helix (F289 vs. Y279) and the intervening linker motif (H285 vs. Q275). Serine gatekeeper residues are uniquely found in ALK4/5/7 offering a strategy for the design of kinome wide selectivity.

**Figure 2.**
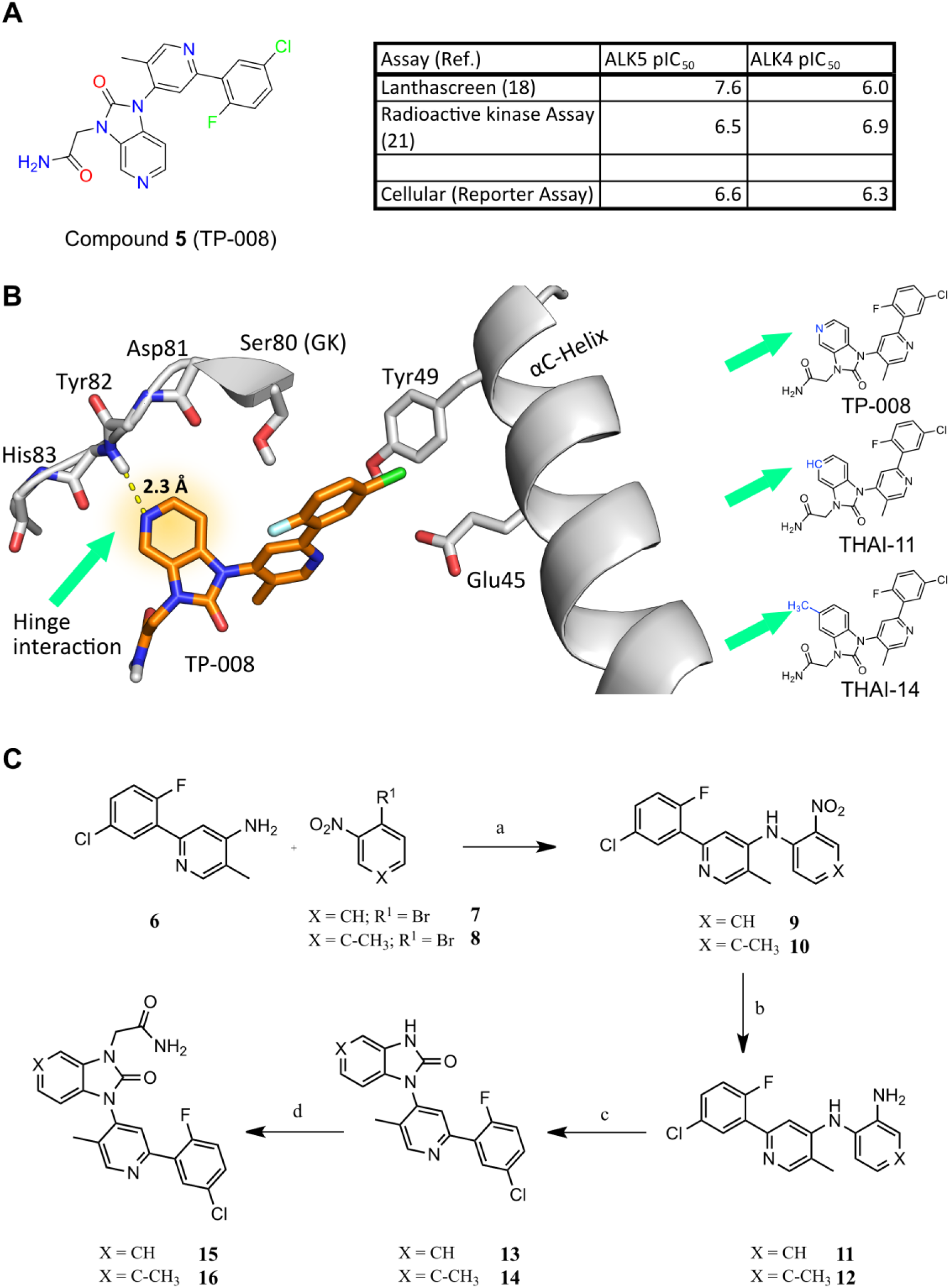
(A) Biochemical and cellular potency of TP-008 on ALK4 and 5, respectively (**5**). (B) Molecular modeling of TP-008 (**5**) in ALK5 and design strategy for the negative control compound. (C) Synthesis route for the potential negative control compounds THAI11 (**15**) and THAI14 (**16**): a) Pd(OAc)_2_, BINAP, Cs_2_CO_3_, toluene, 95 °C, 18 h; b) Fe, HOAc, 100 °C, 30 min; c) CDI, THF, RT, 18 h; d) 2-bromoacetamide, Cs_2_CO_3_, DMF, 65 °C, 3 h.

**Figure 3.**
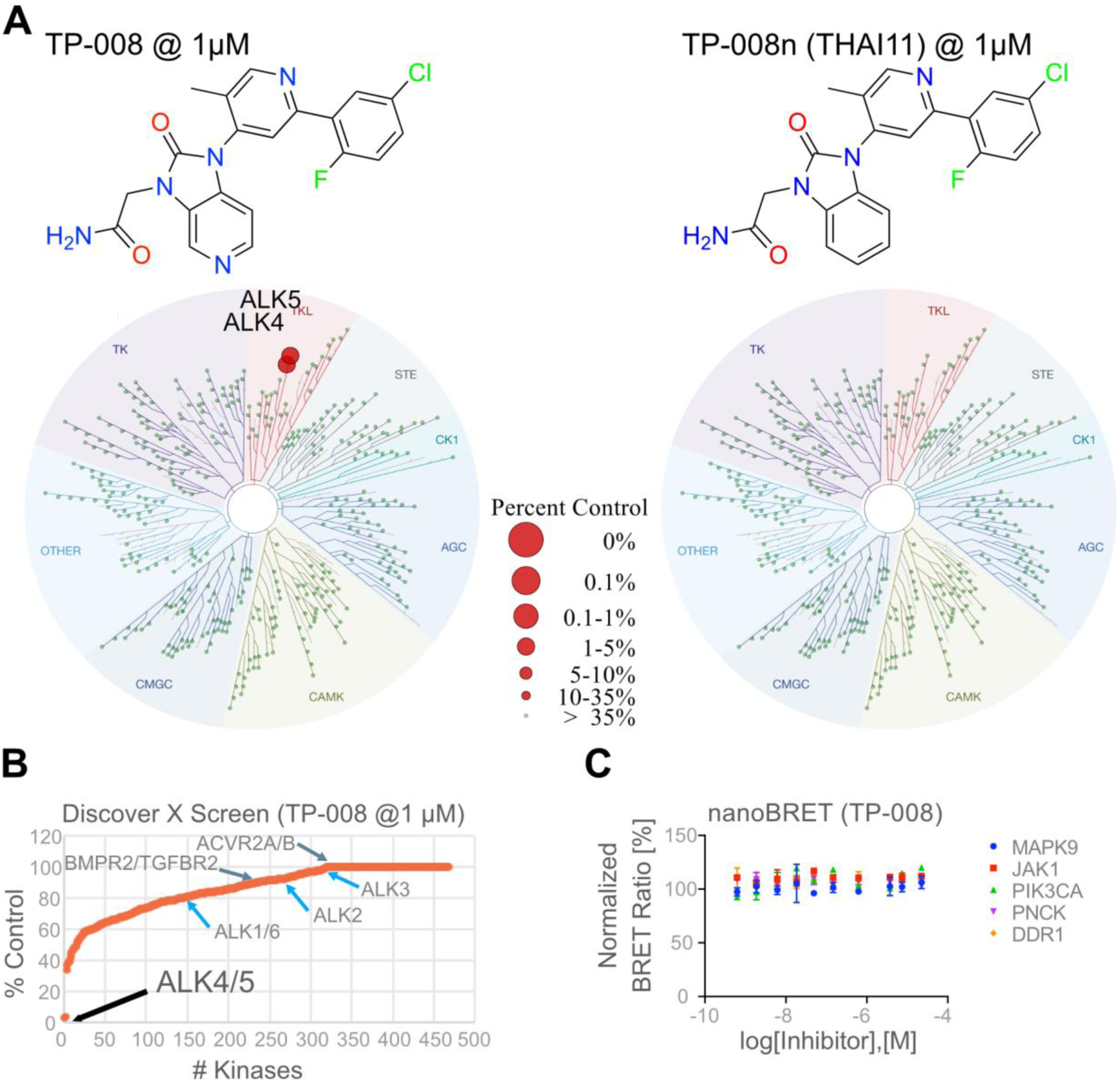
(A) KINOMEscan® of TP-008 (**5**) and THAI11 (**15**) at 1 µM against 469 kinases (*scan*MAX^SM^). Top hits are highlighted as red dots. (B) Waterfall plot, showing selectivity of TP-008 (**5**) against other members from the TGF-β type II receptor and type I receptor family and top hits of TP-008 (**5**) with percent control below 50 % derived from the *scan*Max^SM^ Kinase Assay Panel by Eurofins (DiscoverX). (C) Lack of activity of TP-008 (**5**), THAI11 (**15**) and THAI14 (**16**) against the next top hits from the *scan*MAX^SM^ Kinase Assay, determined by NanoBRET™.

Next, we performed molecular docking studies with TP-008 (**5**) using the ALK5 structure (PDB-ID: 3HMM *(22)*) in order to understand the possible binding mode and as a basis for the design of a related negative control. In all binding modes generated, the pyridine nitrogen of TP-008 (**5**) mediated the interaction with the hinge region (Figure 2B). Therefore, two closely related derivatives were designed and synthesized, either without the nitrogen or with an additional methyl group to block the interaction with the hinge region (Figure 2C). Synthesis of these compounds was performed in four-steps, starting with the building block 2-(5-chloro-2-fluorophenyl)-5-methylpyridin-4-amine (**6**). **6** was coupled to either 1-bromo-2-nitrobenzene (**7**) or 1-bromo-4-methyl-2-nitrobenzene (**8**) in a Buchwald-Hartwig amination to provide the diarylamines **9** and **10**. The nitro group in **9** and **10** was reduced to the primary amines **11** and **12**. The ring closure of **11** and **12** using CDI afforded the benzimidazolones derivatives **13** and **14**, which were finally coupled to 2-bromoacetamide resulting in two potential negative control compounds **15** (THAI11) and **16** (THAI14). Both compounds were also screened in the *scan*Max^SM^ Kinase Assay Panel by Eurofins (DiscoverX) (Figure 3). For **15**, no significant hits were observed at 1 µM, whereas for **16** NEK5 appeared as an off-target, which was likely to be a false positive hit (s. TableS7). However, we thus chose **15** as negative control compound for TP-008 (**5**).

We next used a cellular NanoBRET™ assay to evaluate the cellular selectivity of compounds **5** (TP-008), **15** (THAI11) and **16** (THAI14) against the closest off-targets of **5** suggested by the *in vitro* kinase assay for compound **5** (namely MAPK9, JAK1, PIK3CA, PNCK, DDR1; Figure. 3B) *(23)*. This assay employs the full length protein kinases fused to a luciferase (NLuc) and is performed in live HEK293T cells under physiological ATP concentrations. We recently showed that it provides the advantage of determining differing inhibition profiles comparing the cellular context to *in vitro* assays *(23)*. All compounds tested in the NanoBRET™ assay had an *IC*_50_ higher than 20 µM for the suggested off-target kinases (Figure 3C), indicating that the activity of **5, 15** and **16** was negligent against these kinases in cells at concentrations relevant to ALK4/5 inhibition.

We assessed the cellular potency of compounds **5, 15** and **16** against their target receptor signaling pathways, employing a SMAD2/3-responsive luciferase-based transcriptional reporter assay, in HEK293 cells stimulated with either TGF-β for ALK5 or activin A for ALK4, respectively. Vactosertib (**1**) and GW788388 (**4**) were tested in parallel for reference (Figure 4A, S3). All three compounds displayed dose-dependent inhibition of the reporter in agreement with their activities in enzyme catalytic assays (Figure 4A, S3). Compound **5** yielded *IC*_50_-values of 245 nM and 526 nM against ALK5 and ALK4, respectively, while Vactosertib was more potent with *IC*_50_-values 24 and 12 nM against ALK5 and ALK4. GW788388 exhibited some selectivity for ALK4 with an *IC*_50_ of 44 nM compared to 454 nM against ALK5. Pleasingly, both negative control compounds **15** and **16**, were inactive up to a compound concentration of 10 µM, validating the *in silico*-based design. In addition to the reporter assay, immunoblotting was performed to study the direct effect of the compounds on phosphorylation of the downstream targets SMAD2/3 in C2C12 cells (Figure 4B). TP-008 (**5**), Vactosertib (**1**) and GW788388 (**4**) inhibited the phosphorylation of SMAD2/3 at 1 µM, whereas Vactosertib (**1**) was already active at 100 nM, which is in good agreement with the promoter gene assay. As expected, the two negative control compounds **15** (THAI11) and **16** (THAI14) did not inhibit phosphorylation of SMAD2/3 at concentrations up to 10 µM.

**Figure 4.**
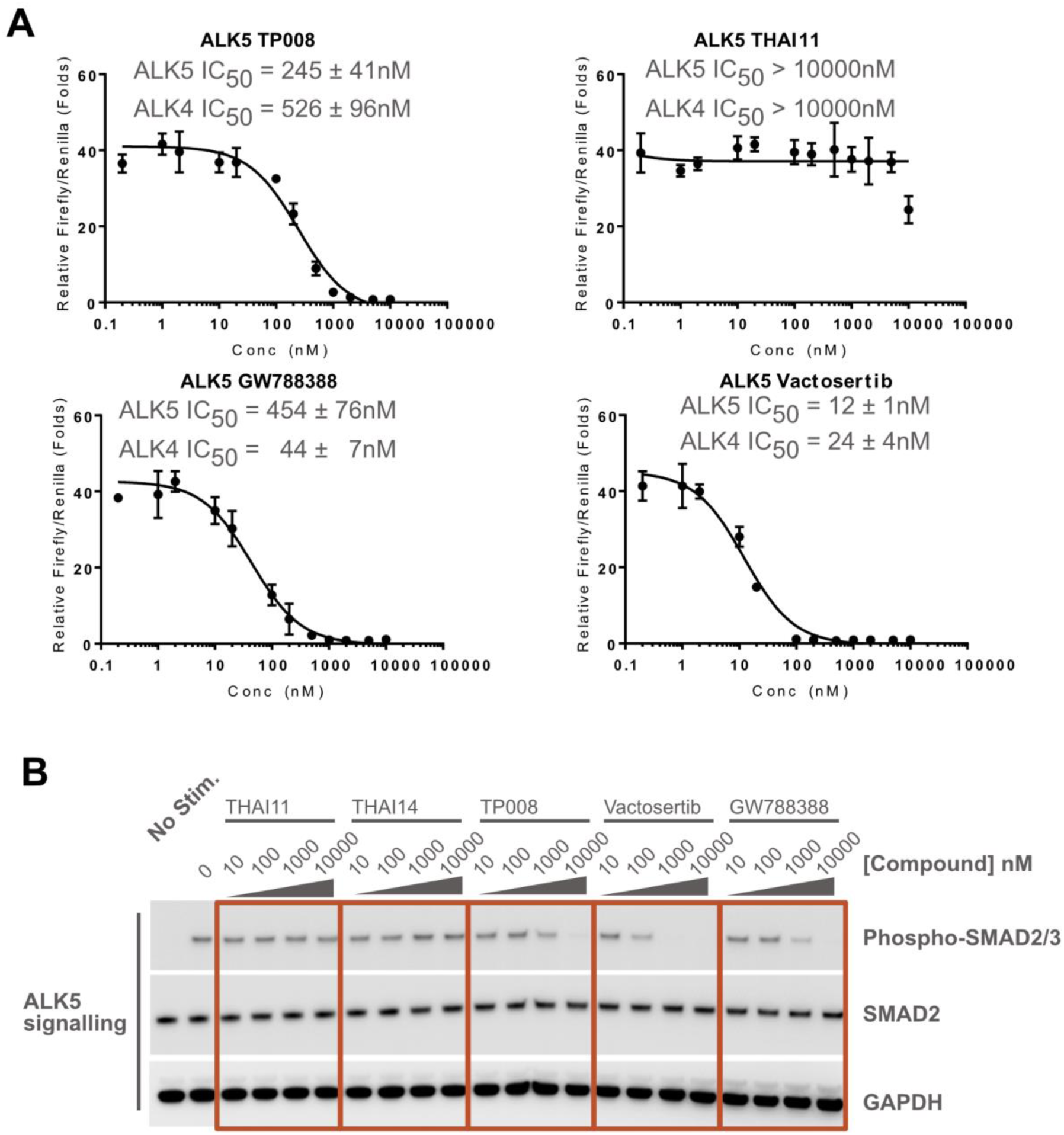
(A) Cellular activity of TP-008 (**5**), THAI11 (**15**), THAI14 (**16**), Vactosertib (**1**) and GW788388 (**4**) on ALK4 and ALK5 measured by a dual luciferase assay using Activin A and TGF-β stimulation, respectively, in HEK293 cells. (B) Immunoblotting showing the inhibitory activity of TP-008 (**5**), THAI11 (**15**), THAI14 (**16**), Vactosertib (**1**) and GW788388 (**4**) on SMAD2/3 phosphorylation in C2C12 cells with or without TGF-β stimulation.

In summary, in this short report we have comprehensively characterized a chemical probe, TP-008, for ALK4/5 and designed two highly related negative control compounds based on a docking model. Many potent ALK5 inhibitors have been published, but their utility for cell signaling studies has been limited by their lack of selectivity. With TP-008, we present a dual inhibitor of ALK4/5 with good cellular potency and the best selectivity profile of available inhibitors so far. Molecular docking indicates that selectivity is driven by the 2-fluoro-5-chlorophenyl moiety, which penetrates deep into the hydrophobic back pocket to pack between the α-C helix residues Tyr49 and Glu45. In addition, ALK4/5 possess a smaller serine residue as the gatekeeper, which is rare within the human kinome. The generated negative control compounds were inactive in the complete KINOMEscan® as well as in the cellular assays. However, we would like to emphasize that the closest paralog of ALK4/5, which is ALK7 as described above in the sequence alignment, was neither part of the *scan*MAX^SM^ Panel by Eurofins/DiscoverX, nor a functional assay that is commercial available by Reaction Biology. Therefore, we cannot exclude activity of TP-008 on this kinase. To conclude, with TP-008 and THAI11, which we henceforth rename as TP-008n, we can present a chemical probe package consisting of a positive and negative control compounds that display the best selectivity of all known ALK5 inhibitors. Importantly, the probe compound TP-008 was able to inhibit the phosphorylation of downstream substrates, whereas TP-008n showed no effect. We recommend using both compounds at a concentration of 1 µM to study the effects of inhibiting ALK5 and ALK4 in cells.

## METHODS

### Compounds and Chemistry

The structures of the presented compounds were verified by ^1^H, ^13^C NMR, and mass spectrometry (ESI); the purity (>95%) of the final compounds was determined by HPLC plus additional high resolution mass spectrometry (HRMS). Compound **1** and **4** were purchased from Selleckchem (München, Germany), while compound **5** and building block **6**, were provided by Takeda. All commercial chemicals and solvents were of reagent grade and were used without further purification. ^1^H and ^13^C NMR spectra were measured in DMSO-d_6_ on a Bruker DPX 250 or AV500 spectrometer. Chemical shifts are reported in parts per million (ppm) in the scale relative to DMSO-d_6_, 2.50 ppm for ^1^H NMR and 39.52 for ^13^C NMR. Mass spectra were obtained on two ThermoFisher Surveyor MSQ’s, (one MSQ coupled to a Camag TLC-MS interface 2 for direct measurements of TLC spots) measuring in the positive- and/or negative-ion mode. High resolution mass spectra were recorded on a MALDI LTQ ORBITRAP XL instrument (Thermo Fisher Scientific). Compound purity was analyzed on a LCMS 2020 from Shimadzu (Duisburg, Germany), under the use of a MultoHigh 100 RP 18, 3 μ, 100 × 2 mm column from CS Chromatography-Service GmbH (Langerwehe, Germany) with an acetonitrile/water gradient from 20–75% at a flow rate of 1 mL/min and UV detection at 245 and 280 nm.

### Cell culture

HEK-293 and C2C12 cells were maintained in DMEM medium (Gibco) supplemented with 10% fetal bovine serum (FBS) (Thermo Fisher), and 10 µg/mL gentamicin (Sigma).

### HEK-293 transfection and dual luciferase reporter assay

CAGA-Luc and Renilla-luciferase constructs (a gift of Dr Petra Knaus, Free University of Berlin) were used as the reporter for ALK4/ALK5 signalling and loading control respectively. HEK-293 cells were transfected with reporter constructs using FuGENE HD (Promega) according to the manufacturer’s instructions. Briefly, 4 parts of CAGA-Luc construct was mixed with 1 part of Renilla-luciferase construct (mass/mass) and diluted into phenol red-free Opti-MEM (Gibco) at a concentration of 10 µg/mL. Without coming in contact with the sides of the container, 3µL FuGENE HD was added for each µg of DNA used. After thorough mixing by inversion, FuGENE HD/DNA complexes were allowed to form by incubation at room temperature for 20 minutes. 1 part of transfection mixture was added to 20 parts of HEK-293 cell suspension with a density of 200,000 cells per mL (volume/volume). 50 µL of HEK-293 cell suspension (10,000 cells) was dispensed into each well of 96-well plate (Corning) and incubated in a humidified, 37 °C incubator with 5% carbon dioxide. 24 hours after transfection, the cells were incubated with ligand and test compounds (**1**; **4**; **5**; **15** and **16**) diluted in 2 mL of DMEM medium with 1% fetal bovine serum at the indicated concentrations. 100 ng/mL Activin A (120-14-100, Peprotech) and 10 ng/mL TGF-β (100-21-10, Peptrotech) was used for ALK4 and ALK5 stimulation respectively. After 24 hours, cells were harvested, lysed and processed for measurement of luciferase activity using the Dual-Luciferase Reporter Assay System (Promega) according to the manufacturer’s instructions. Briefly, culture medium was aspirated completely and cells were lysed in 50 µL of 1X PLB with 300 rpm agitation for 30 minutes. 10 µL of cell lysate was dispensed into each well of 384-well flat bottom polypropylene plate (Greiner). Luminescent signal of firefly and Renilla luciferase activity were measured sequentially using PHERAstar FS microplate reader (BMG Labtech) after the addition of 25 µL of LARII and Stop & Glo respectively. A measurement interval of 2 seconds and gain setting of 3600 were used. Firefly luciferase signal was normalised to cell number by division with Renilla luciferase signal. Relative luciferase unit (RLU) was obtained by further division with signal from cells without ligand stimulation. The apparent EC_50_ values of test compounds (**1**; **4**; **5**; **15** and **16**) were estimated using the [Iinhibitor] vs. response (three parameters) non-linear regression curve fitting function of GraphPad Prism 7.

### Western blot analysis

500,000 C2C12 cells in 2 mL of culture medium were seeded into each well of 6-well plates (Corning) and incubated overnight in a humidified, 37 °C incubator with 5% carbon dioxide to allow cell attachment. Cells were serum-starved for 1 hour in 2 mL of Opti-MEM (Gibco). Cells were subsequently incubated with ligand and test compounds (**1**; **4**; **5**; **15** and **16**) diluted in 1 mL of Opti-MEM at the indicated concentrations. After 1 hour, Opti-MEM was aspirated completely and 400 µL of lysis buffer (150 mM NaCl, 20 mM Tris-HCl pH 7.5, 1 % Triton-X100, supplemented with complete mini EDTA-free protease inhibitor cocktail (Sigma), 25 mM NaF and 25 mM Na_3_VO_4_) was added to each well. 6-well plates were agitated at 400 rpm for 20 minutes at 4 °C for the complete lysis of cells. Cell lysates were transferred into 1.5 mL tubes and centrifuged at 15000 XG for 10 minutes at 4 °C. Supernatant was collected and mixed with sample loading buffer (62.5 mM Tris-HCl pH 6.8, 2.5 % SDS, 0.002 % bromophenol blue, 5 % β-mercaptoethanol, 10 % glycerol). Samples were boiled for 5 minutes at 95 °C. 10 µL of samples were loaded per lane for SDS-PAGE. After gel transfer, PVDF membrane was blocked in PBS-T (137 mM NaCl, 27 mM KCl, 100 mM Na_2_PO_4_, 18 mM KH_2_PO_4_, 0.1 % Tween-20) supplemented with 3 % bovine serum albumin for 1 hour at room temperature. PVDF membrane was incubated with primary antibody overnight at 4 °C with agitation. Anti-phospho SMAD2/3 (8828S, Cell Signalling Technology) and anti-SMAD2 (5339S, Cell Signalling Technology) were diluted 1000X and anti-GAPDH (AM4300, Thermo Fisher) was diluted 5000X in PBST-T supplemented with 3 % bovine serum albumin. After washing 3 times for 15 minutes with PBS-T, PVDF membrane was incubated with secondary antibody-HRP for 1 hour at room temperature with agitation. Anti-rabbit-HRP (A6667, Sigma) was diluted 1000X and anti-mouse-HRP (GTX213111-01, GeneTex) was diluted 2000X in PBS-T supplemented with 3% skim milk. After washing 3 times for 15 minutes with PBS-T, PVDF membrane was incubated for 1 minute with Pierce ECL Western Blotting Substrate (Thermo Fisher). Luminescent signal on PVDF membrane was imaged using ImageQuant LAS-4000 (GE Healthcare).

### NanoBRET™ target engagement assays

The assay was performed as described previously *(23)*. In brief: Full-length kinase ORF (Promega) cloned in frame with a NanoLuc-vector (as indicated in Table S8) was transfected into HEK293T cells using FuGENE HD (Promega, E2312) and proteins were allowed to express for 20 h. Serially diluted inhibitor and NanoBRET™ Kinase Tracer (as indicated in Table S8) were pipetted into white 384-well plates (Greiner 781 207) using an ECHO 555 acoustic dispenser (Labcyte). The corresponding transfected cells were added and reseeded at a density of 2 × 105 cells/mL after trypsinization and resuspension in Opti-MEM without phenol red (Life Technologies). The system was allowed to equilibrate for 2 hours at 37 °C and 5 % CO_2_ prior to BRET measurements. To measure BRET, NanoBRET™ NanoGlo Substrate + Extracellular NanoLuc Inhibitor (Promega, N2160) were added as per the manufacturer’s protocol, and filtered luminescence was measured on a PHERAstar plate reader (BMG Labtech) equipped with a luminescence filter pair (450 nm BP filter (donor) and 610 nm LP filter (acceptor)). Competitive displacement data were then plotted using GraphPad Prism 8 software using a 4-parameter curve fit with the following equation: Y=Bottom + (Top-Bottom) / (1+10^((LogIC50-X)*HillSlope))

### Docking-Study

Molecular docking studies on ALK5 were performed with the platform seeSAR v8.1 *(24)*. The crystal structure 3HHM was selected due to similar structural features between the co-crystallized ligand and TP-008. The binding site was defined using the co-crystallized ligand as the center point for selecting amino acids in the radius of 6.5 Å. Re-docking of the co-crystallized ligand showed similar binding pose as the reference, presenting the same interactions with the hinge region and Lys 32 of ALK5. Following this analysis, TP-008 was docked and 10 poses were generated. All 10 poses showed very similar binding modes, with the pyridine moiety interacting with the hinge region, while the main differences observed were from the flexibility of the amide moiety, with possible interactions with the backbone of residues Ile11 and Ser87 and hydrogen bond with residue Asp90.

## ASSOCIATED CONTENT

### Supporting Information Available

This material is available free of charge via the Internet.

## ACKNOWLEDGEMENTS

J.F.W. acknowledges support from The Brain Tumour Charity. The SGC is a registered charity (number 1097737) that receives funds from AbbVie, Bayer Pharma AG, Boehringer Ingelheim, Canada Foundation for Innovation, Eshelman Institute for Innovation, Genome Canada, Innovative Medicines Initiative (EU/EFPIA) [ULTRA-DD grant no. 115766], Janssen, Merck KGaA Darmstadt Germany, MSD, Novartis Pharma AG, Ontario Ministry of Economic Development and Innovation, Pfizer, São Paulo Research Foundation-FAPESP, Takeda, and Wellcome [106169/ZZ14/Z].

## For Table of Contents Only

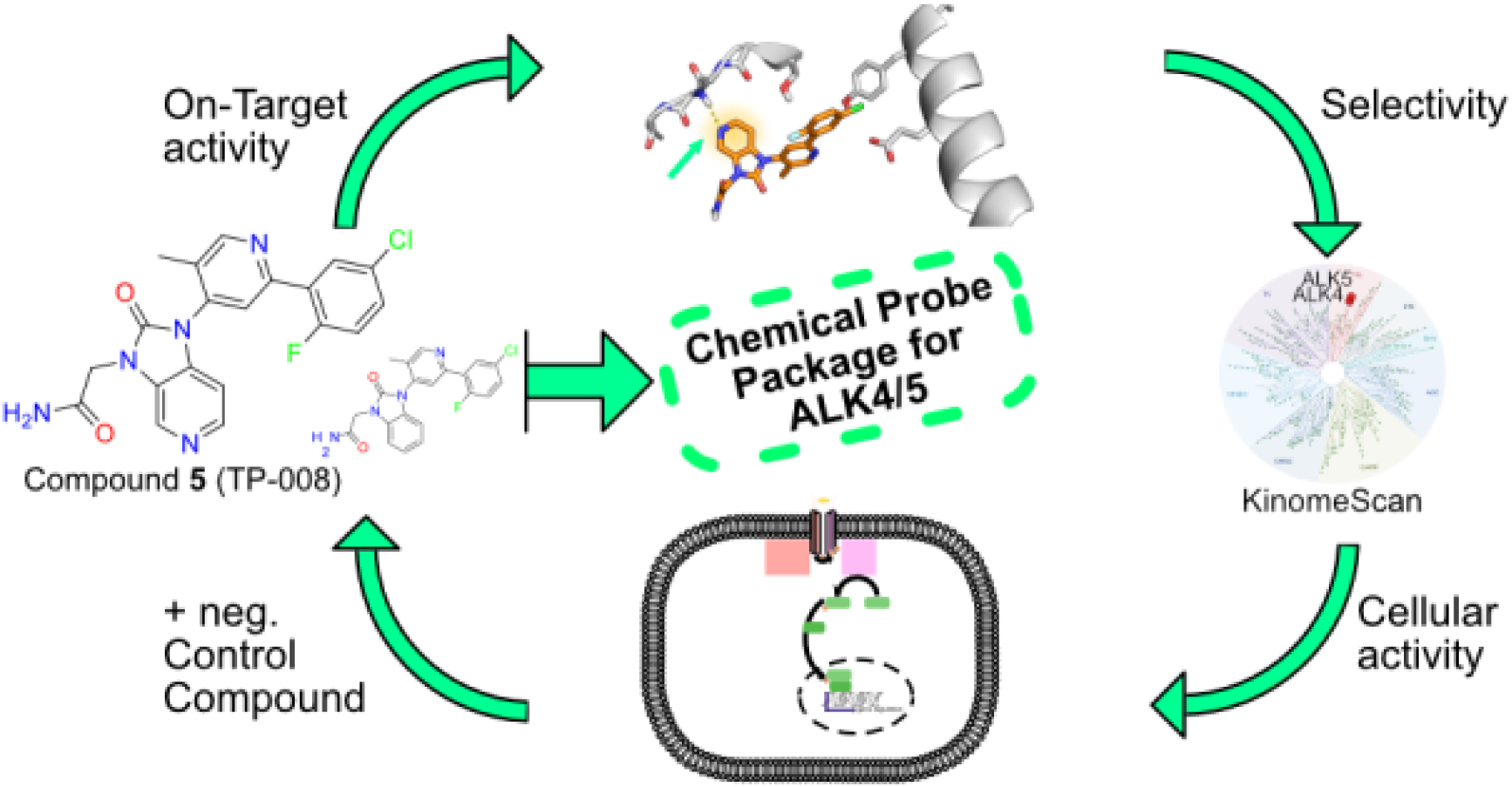

