## Supporting info for "A Highly Selective Chemical Probe for Activin Receptor-like Kinases ALK4 and ALK5"

Thomas Hanke[1], Jong Fu Wong[2], Benedict-Tilmann Berger[1], Ismahan Abdi[1], Lena Marie Berger[1], Roberta Tesch[1], Claudia Tredup[1], Alex Bullock[2], Susanne Müller[1], Stefan Knapp[1]
[1]Institute for Pharmaceutical Chemistry and Buchmann Institute for Molecular Life Sciences, Johann Wolfgang Goethe-University, Max-von-Laue-Str. 9, D-60438 Frankfurt am Main, Germany.
[2]Nuffield Department of Clinical Medicine, Structural Genomics Consortium, University of Oxford, NDM Research Building, Roosevelt Drive, Oxford, OX3 7FZ, UK.

**– Supplementary Information –**

### I Supplementary Figures and Tables

**Table S1.** $IC_{50}$ summary of enzymatic kinase assay.

| | Compound $IC_{50}$* (M): | | | | $IC_{50}$ (M) Control Cmpd | Control Cmpd ID |
|---|---|---|---|---|---|---|
| **Kinase:** | **TP-008** | **THAI11** | **GW788388** | **VACTOSERTIB** | | |
| **ALK4/ ACVR1B** | 1.13E-07 | >1.00E-05 | 1.55E-07 | 8.30E-09 | 8.86E-08 | **LDN193189** |
| **ALK5/ TGFBR1** | 3.43E-07 | >1.00E-05 | 2.22E-07 | 1.18E-08 | 2.36E-07 | **LDN193189** |

**Figure S1.** Compound $IC_{50}$ Data of compound (**1**), (**4**), (**5**) and (**16**) for ALK4.

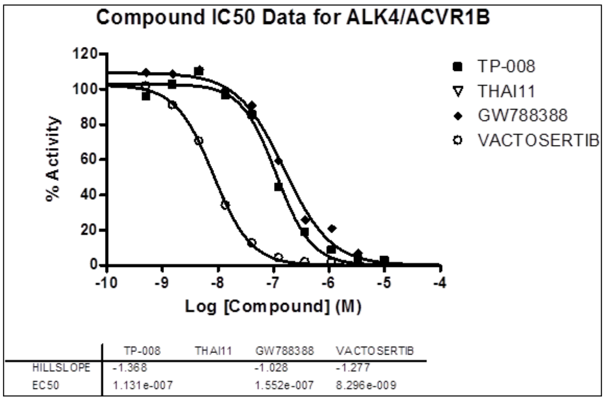

Assay was performed as described in chapter II) Radioactive kinase Assay from reaction biology.

**Figure S2.** Compound $IC_{50}$ Data of compound (**1**), (**4**), (**5**) and (**16**) for ALK5.

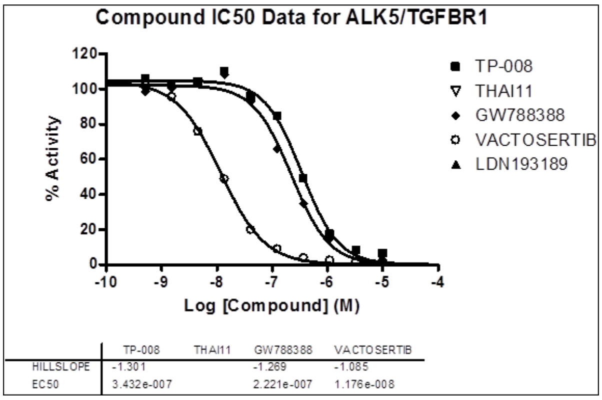

Assay was performed as described in chapter II) Radioactive kinase Assay from reaction biology.

**Figure S3.** Cellular activity of THAI14 (**16**) on ALK4/5, determined by a dual luciferase assay.

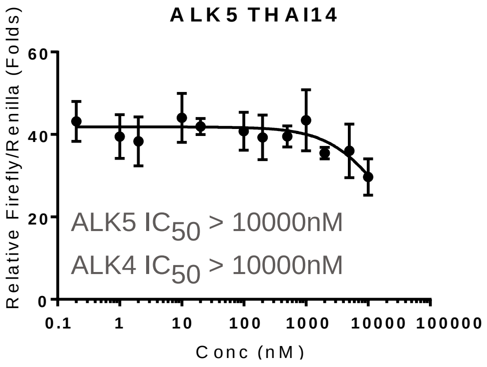

**Table S2.** Sequence Alignment of ALK1–7 of key residues in the hinge-region.

| Kinase Residue | g.loop | | | | | | αC-Helix | | | | | | | | | | | GK | Hinge | | | linker | | | | αD-Helix | | | | | | | αE-Helix | | | | | | xDFG | | | |
|---|---|---|---|---|---|---|---|---|---|---|---|---|---|---|---|---|---|---|---|---|---|---|---|---|---|---|---|---|---|---|---|---|---|---|---|---|---|---|---|---|---|---|
| ALK1 | G | K | G | R | Y | G | S | W | F | R | E | T | E | I | Y | N | T | T | H | Y | H | E | H | G | S | L | Y | D | F | L | Q | R | T | Q | G | K | P | A | A | D | L | G |
| ALK2 | G | K | G | R | Y | G | S | W | F | R | E | T | E | L | Y | N | T | T | H | Y | H | E | M | G | S | L | Y | D | Y | L | Q | L | T | Q | G | K | P | A | A | D | L | G |
| ALK3 | G | K | G | R | Y | G | S | W | F | R | E | T | E | I | Y | Q | T | T | D | Y | H | E | N | G | S | L | Y | D | F | L | K | C | T | Q | G | K | P | A | A | D | L | G |
| ALK4 | G | K | G | R | F | G | S | W | F | R | E | A | E | I | Y | Q | T | S | D | Y | H | E | H | G | S | L | F | D | Y | L | N | R | T | Q | G | K | P | G | A | D | L | G |
| ALK5 | G | K | G | R | F | G | S | W | F | R | E | A | E | I | Y | Q | T | S | D | Y | H | E | H | G | S | L | F | D | Y | L | N | R | T | Q | G | K | P | A | A | D | L | G |
| ALK6 | G | K | G | R | Y | G | S | W | F | R | E | T | E | I | Y | Q | T | T | D | Y | H | E | N | G | S | L | Y | D | Y | L | K | S | T | Q | G | K | P | A | A | D | L | G |
| ALK7 | G | K | G | R | F | G | S | W | F | R | E | A | E | I | Y | Q | T | S | E | Y | H | E | Q | G | S | L | Y | D | Y | L | N | R | T | Q | G | K | P | A | A | D | L | G |

**Table S3.** Sequence Alignment of ALK4/5.

```
ALK-4          13 LVVLLLAGSGGSGPRGVQALLCACTSCLQANYTCETDGACMVSIFNLDGMEHHVRTCIPK
ALK-5          15 LVLAAAAAAAAALLPGATALQCFCHLCTKDNFTCVTDGLCFVSVTETTDKVIHNSMCIAE
                  **    *        *  ** * *  *   * ** *** * **         *   **

ALK-4          73 VELVPAGKPFYCLSSE---DLRNTHCCYTDYCNRIDLRVPSGHLKEPEHPSMWGPVELVG
ALK-5          75 IDLIPRDRPFVCAPSSKTGSVTTTYCCNQDHCNKIEL--PTTVKSSPG----LGPVELAA
                    * *   ** *  *        * **  * ** * *  *      *      *****

ALK-4         130 IIAGPVFLLFLIIIIVFLVINYHQRVYHNRQRLDMEDPSCEM-CLSKDKTLQDLVYDLST
ALK-5         129 VIAGPV--CFVCISLMLMVYICHNRTVIHHRVPNEEDPSLDRPFISEGTTLKDLIYDMTT
                   *****   *  *     *   * *          ****      *   ** ** **  *
                                    Beta-1 P-LoopBeta-2   Beta-3     alpha-C
ALK-4         189 SGSGSGLPLFVQRTVARTIVLQEIIGKGRFGEVWRGRWRGGDVAVKIFSSREERSWFREA
ALK-5         187 SGSGSGLPLLVQRTIARTIVLQESIGKGRFGEVWRGKWRGEEVAVKIFSSREERSWFREA
                  ********* **** ******** ************ ***  ******************
                                   Beta-4     Beta-5 Hinge alpha-D   alpha-E
ALK-4         249 EIYQTVMLRHENILGFIAADNKDNGTWTQLWLVSDYHEHGSLFDYLNRYTVTIEGMIKLA
ALK-5         247 EIYQTVMLRHENILGFIAADNKDNGTWTQLWLVSDYHEHGSLFDYLNRYTVTVEGMIKLA
                  **************************************************** *******
                                      Beta-6    Beta-7   Beta-8   Beta-9
ALK-4         309 LSAASGLAHLHMEIVGTQGKPGIAHRDLKSKNILVKKNGMCAIADLGLAVRHDAVTDTID
ALK-5         307 LSTASGLAHLHMEIVGTQGKPAIAHRDLKSKNILVKKNGTCCIADLGLAVRHDSATDTID
                  ** ****************** ***************** * ***********  *****

ALK-4         369 IAPNQRVGTKRYMAPEVLDETINMKHFDSFKCADIYALGLVYWEIARRCNSGGVHEEYQL
ALK-5         367 IAPNHRVGTKRYMAPEVLDDSINMKHFESFKRADIYAMGLVFWEIARRCSIGGIHEDYQL
                  **** **************  ****** *** ***** *** *******  ** ** ***

ALK-4         429 PYYDLVPSDPSIEEMRKVVCDQKLRPNIPNWWQSYEALRVMGKMMRECWYANGAARLTAL
ALK-5         427 PYYDLVPSDPSVEEMRKVVCEQKLRPNIPNRWQSCEALRVMAKIMRECWYANGAARLTAL
                  *********** ******** ********* *** ****** * ****************
```

```
ALK-4        489 RIKKTLSQLSVQEDVKI
ALK-5        487 RIKKTLSQLSQQEGIKM
                 ********** **  *
```

**Table S4.** Sequence Alignment of ALK1–7.

```
CLUSTAL O(1.2.4) multiple sequence alignment

SP|P37023|ACVL1_HUMAN       ALK1      -------------------MTLGSPRKGL-LML-------LMALVTQGDPVKPSRGPLVT 33
SP|Q04771|ACVR1_HUMAN       ALK2      -------------------MVDGVMILPVLIMI-------ALPSPSMEDEKPKVNPKLYM 34
SP|P36894|BMR1A_HUMAN       ALK3      MPQLYIYIRLLGAYLFIISRVQGQNLDSMLHGTGMKSDSDQKKSENGVTLAPEDTLPFLK 60
SP|P36896|ACV1B_HUMAN       ALK4      -----------------MAES--AGASSFFPL--------VVLLLAGSGGSGPRGVQALL 33
SP|P36897|TGFR1_HUMAN       ALK5      -----------------MEAAVAAPRPRLLLL--------VLAAAAAAAALLPGATALQ 35
SP|O00238|BMR1B_HUMAN       ALK6      ---------------------------MLLRSAGKL--NVGTKKEDGESTAPTPRPKVLR 31
SP|Q8NER5|ACV1C_HUMAN       ALK7      -----------------MTRALCSALRQ------------ALLLL-AAAAEL---SPGLK 27

SP|P37023|ACVL1_HUMAN       ALK1      CTCESP---HCKG-PTCRGAWCTVVLVREEGRHPQEHRGCGNL-------HRELCRGRPT 82
SP|Q04771|ACVR1_HUMAN       ALK2      CVCEGL---SCGNEDHCEGQQCFSSLSINDGFHVY-QKGCFQV----YEQGKMTCKTPPS 86
SP|P36894|BMR1A_HUMAN       ALK3      CYCSGHCPDDAINNTCITNGHCFAIIEEDDQGETTLASGCMK-----YEGSDFQCKDSPK 115
SP|P36896|ACV1B_HUMAN       ALK4      CACTSC---LQANYTCETDGACMVSIFNLDGMEHHV-RTCIPKVELVPAGKPFYCLSSED 89
SP|P36897|TGFR1_HUMAN       ALK5      CFCHLC---TKDNFTCVTDGLCFVSVTETTDKVIHN-SMCIAEIDLIPRDRPFVCAPSSK 91
SP|O00238|BMR1B_HUMAN       ALK6      CKCHHHCPEDSVNNICSTDGYCFTMIEEDDSGLPVVTSGCLG-----LEGSDFQCRDTPI 86
SP|Q8NER5|ACV1C_HUMAN       ALK7      CVCLLC---DSSNFTCQTEGACWASVMLTNGKEQVI-KSCVSLPEL---NAQVFCHSSNN 80
                                      * *         .        *   :             *              *

SP|P37023|ACVL1_HUMAN       ALK1      E--FVNHYCCD-SHLCNHNVSLVLEATQPPSEQPGTDGQ---LAL-ILGPVLALLALVAL 135
SP|Q04771|ACVR1_HUMAN       ALK2      P--GQAVECCQ-GDWCNRNITAQLPTKGK--SFPGTQNFHLEVGLIILSVVFAVCLLACL 141
SP|P36894|BMR1A_HUMAN       ALK3      AQLRRTIECCR-TNLCNQYLQPT-LPPVVIG--PFFDGSIRWLVLLISMAVCIIAMIIFS 171
SP|P36896|ACV1B_HUMAN       ALK4      ---LRNTHCCY-TDYCNRIDLRVPSGHLKEPEHPSMWGPV-ELVGIIAGPVFLLFLIIII 144
SP|P36897|TGFR1_HUMAN       ALK5      TGSVTTTYCCN-QDHCNKIELPTTV------KSSPGLGPV-ELAAVIAGPVCFVCISLML 143
SP|O00238|BMR1B_HUMAN       ALK6      PHQRRSIECCTERNECNKDLHPT-LPPLKNR--DFVDGPIHHRALLISVTVCSLLLVL-I 142
SP|Q8NER5|ACV1C_HUMAN       ALK7      ---VTKTECCF-TDFCNNITLHLPTAS----PNAPKLGPM-ELAIIITVPVCLLSIAAML 131
                                    **   . **.                    .        *   *  :

SP|P37023|ACVL1_HUMAN       ALK1      GVLGLWHV-RRRQEKQRGLHSELGESSLILKASEQGDSMLGDLLDSDCTTGSGSGLPFLV 194
SP|Q04771|ACVR1_HUMAN       ALK2      LGVALRKFKRRNQERLNPRDVEYGTIE-GLITTNVGDSTLADLLDHSCTSGSGSGLPFLV 200
SP|P36894|BMR1A_HUMAN       ALK3      SCFCYKHYCKSISSRRR-YNRDLEQDE----AFIPVGESLKDLIDQSQSSGSGSGLPLLV 226
SP|P36896|ACV1B_HUMAN       ALK4      VFLVINYHQRVYHNRQRLDMEDPSC-E----MCLSKDKTLQDLVYDLSTSGSGSGLPLFV 199
SP|P36897|TGFR1_HUMAN       ALK5      MVYIC-HNRTVIHHRV-PNEEDPSLDR----PFISEGTTLKDLIYDMTTSGSGSGLPLLV 197
SP|O00238|BMR1B_HUMAN       ALK6      ILFCYFRYKR-QETRPR-YSIGLEQDE----TYIPPGESLRDLIEQSQSSGSGSGLPLLV 196
SP|Q8NER5|ACV1C_HUMAN       ALK7      TVWACQGRQCSYRKKKRPNVEEPLSEC----NLVNAGKTLKDLIYDVTASGSGSGLPLLV 187
                                                    :                     .  * **:    ::*******::*

                                                      g.l                          αC
SP|P37023|ACVL1_HUMAN       ALK1      QRTVARQVALVECVGKGRYGEVWRGLWHGESVAVKIFSSRDEQSWFRETEIYNTVLLRHD 254
SP|Q04771|ACVR1_HUMAN       ALK2      QRTVARQITLLECVGKGRYGEVWRGSWQGENVAVKIFSSRDEKSWFRETELYNTVMLRHE 260
SP|P36894|BMR1A_HUMAN       ALK3      QRTIAKQIQMVRQVGKGRYGEVWMGKWRGEKVAVKVFFTTEEASWFRETEIYQTVLMRHE 286
SP|P36896|ACV1B_HUMAN       ALK4      QRTVARTIVLQEIIGKGRFGEVWRGRWRGGDVAVKIFSSREERSWFREAEIYQTVMLRHE 259
SP|P36897|TGFR1_HUMAN       ALK5      QRTIARTIVLQESIGKGRFGEVWRGKWRGEEVAVKIFSSREERSWFREAEIYQTVMLRHE 257
SP|O00238|BMR1B_HUMAN       ALK6      QRTIAKQIQMVKQIGKGRYGEVWMGKWRGEKVAVKVFFTTEEASWFRETEIYQTVLMRHE 256
SP|Q8NER5|ACV1C_HUMAN       ALK7      QRTIARTIVLQEIVGKGRFGEVWHGRWCGEDVAVKIFSSRDERSWFREAEIYQTVMLRHE 247
                                      ***:*: : : . :****:**** * * * .****:* : :* *****:*:*:**::**:
                                                          GK/Hinge/linker/αD/
SP|P37023|ACVL1_HUMAN       ALK1      NILGFIASDMTSRNSSTQLWLITHYHEHGSLYDFLQRQTLEPHLALRLAVSAACGLAHLH 314
SP|Q04771|ACVR1_HUMAN       ALK2      NILGFIASDMTSRHSSTQLWLITHYHEMGSLYDYLQLTTLDTVSCLRIVLSIASGLAHLH 320
SP|P36894|BMR1A_HUMAN       ALK3      NILGFIAADIKGTGSWTQLYLITDYHENGSLYDFLKCATLDTRALLKLAYSAACGLCHLH 346
SP|P36896|ACV1B_HUMAN       ALK4      NILGFIAADNKDNGTWTQLWLVSDYHEHGSLFDYLNRYTVTIEGMIKLALSAASGLAHLH 319
SP|P36897|TGFR1_HUMAN       ALK5      NILGFIAADNKDNGTWTQLWLVSDYHEHGSLFDYLNRYTVTVEGMIKLALSTASGLAHLH 317
SP|O00238|BMR1B_HUMAN       ALK6      NILGFIAADIKGTGSWTQLYLITDYHENGSLYDYLKSTTLDAKSMLKLAYSSVSGLCHLH 316
SP|Q8NER5|ACV1C_HUMAN       ALK7      NILGFIAADNKDNGTWTQLWLVSEYHEQGSLYDYLNRNIVTVAGMIKLALSIASGLAHLH 307
                                      *******:* ..  : ***:*::.*** ***:*:*:   :     :::. * ..**.***
                                            αE                        xDFGa.l
SP|P37023|ACVL1_HUMAN       ALK1      VEIFGTQGKPAIAHRDFKSRNVLVKSNLQCCIADLGLAVMHSQGSDYLDIGNNPRVGTKR 374
SP|Q04771|ACVR1_HUMAN       ALK2      IEIFGTQGKPAIAHRDLKSKNILVKKNGQCCIADLGLAVMHSQSTNQLDVGNNPRVGTKR 380
SP|P36894|BMR1A_HUMAN       ALK3      TEIYGTQGKPAIAHRDLKSKNILIKKNGSCCIADLGLAVKFNSDTNEVDVPLNTRVGTKR 406
SP|P36896|ACV1B_HUMAN       ALK4      MEIVGTQGKPGIAHRDLKSKNILVKKNGMCAIADLGLAVRHDAVTDTIDIAPNQRVGTKR 379
SP|P36897|TGFR1_HUMAN       ALK5      MEIVGTQGKPAIAHRDLKSKNILVKKNGTCCIADLGLAVRHDSATDTIDIAPNHRVGTKR 377
SP|O00238|BMR1B_HUMAN       ALK6      TEIFSTQGKPAIAHRDLKSKNILVKKNGTCCIADLGLAVKFISDTNEVDIPPNTRVGTKR 376
SP|Q8NER5|ACV1C_HUMAN       ALK7      MEIVGTQGKPAIAHRDIKSKNILVKKCETCAIADLGLAVKHDSILNTIDIPQNPKVGTKR 367
                                       ** .*****.*****:**:*:*:*.   *.******** .    : :*:  * :*****

SP|P37023|ACVL1_HUMAN       ALK1      YMAPEVLDEQIRTDCFESYKWTDIWAFGLVLWEIARRTIVNGIVEDYRPPFYDVVPNDPS 434
SP|Q04771|ACVR1_HUMAN       ALK2      YMAPEVLDETIQVDCFDSYKRVDIWAFGLVLWEVARRMVSNGIVEDYKPPFYDVVPNDPS 440
SP|P36894|BMR1A_HUMAN       ALK3      YMAPEVLDESLNKNHFQPYIMADIYSFGLIIWEMARRCITGGIVEEYQLPYYNMVPSDPS 466
SP|P36896|ACV1B_HUMAN       ALK4      YMAPEVLDETINMKHFDSFKCADIYALGLVYWEIARRCNSGGVHEEYQLPYYDLVPSDPS 439
SP|P36897|TGFR1_HUMAN       ALK5      YMAPEVLDDSINMKHFESFKRADIYAMGLVFWEIARRCSIGGIHEDYQLPYYDLVPSDPS 437
SP|O00238|BMR1B_HUMAN       ALK6      YMPPEVLDESLNRNHFQSYIMADMYSFGLILWEVARRCVSGGIVEEYQLPYHDLVPSDPS 436
SP|Q8NER5|ACV1C_HUMAN       ALK7      YMAPEMLDDTMNVNIFESFKRADIYSVGLVYWEIARRCSVGGIVEEYQLPYYDMVPSDPS 427
                                      ** **:**: :. . *: :  .*:::.**: **:***   .*: *:*: *::::**.***
```

```
SP|P37023|ACVL1_HUMAN           ALK1      FEDMKKVVCVDQQTPTIPNRLAADPVLSGLAQMMRECWYPNPSARLTALRIKKTLQKISN 494
SP|Q04771|ACVR1_HUMAN           ALK2      FEDMRKVVCVDQQRPNIPNRWFSDPTLTSLAKLMKECWYQNPSARLTALRIKKTLTKIDN 500
SP|P36894|BMR1A_HUMAN           ALK3      YEDMREVVCVKRLRPIVSNRWNSDECLRAVLKLMSECWAHNPASRLTALRIKKTLAKMVE 526
SP|P36896|ACV1B_HUMAN           ALK4      IEEMRKVVCDQKLRPNIPNWWQSYEALRVMGKMMRECWYANGAARLTALRIKKTLSQLSV 499
SP|P36897|TGFR1_HUMAN           ALK5      VEEMRKVVCEQKLRPNIPNRWQSCEALRVMAKIMRECWYANGAARLTALRIKKTLSQLSQ 497
SP|O00238|BMR1B_HUMAN           ALK6      YEDMREIVCIKKLRPSFPNRWSSDECLRQMGKLMTECWAHNPASRLTALRVKKTLAKMSE 496
SP|Q8NER5|ACV1C_HUMAN           ALK7      IEEMRKVVCDQKFRPSIPNQWQSCEALRVMGRIMRECWYANGAARLTALRIKKTISQLCV 487
                                           *:*:::** .:  * . *   :   *  : ::* ***  * ::******:***: ::

SP|P37023|ACVL1_HUMAN           ALK1      SPEKPKVIQ 503
SP|Q04771|ACVR1_HUMAN           ALK2      SLDKLKTDC 509
SP|P36894|BMR1A_HUMAN           ALK3      SQDVKI--- 532
SP|P36896|ACV1B_HUMAN           ALK4      QEDVKI--- 505
SP|P36897|TGFR1_HUMAN           ALK5      QEGIKM--- 503
SP|O00238|BMR1B_HUMAN           ALK6      SQDIKL--- 502
SP|Q8NER5|ACV1C_HUMAN           ALK7      KEDCKA--- 493
                                          .
```

**Table S5**. Kinome scan of TP-008 (**5**) @ 1 µM against 469 kinases from DiscoverX/Eurofins.

| Compound Name | DiscoveRx Gene Symbol | Entrez Gene Symbol | Percent Control |
|---|---|---|---|
| TP-008 | AAK1 | AAK1 | 100 |
| TP-008 | ABL1(E255K)-phosphorylated | ABL1 | 71 |
| TP-008 | ABL1(F317I)-nonphosphorylated | ABL1 | 100 |
| TP-008 | ABL1(F317I)-phosphorylated | ABL1 | 100 |
| TP-008 | ABL1(F317L)-nonphosphorylated | ABL1 | 94 |
| TP-008 | ABL1(F317L)-phosphorylated | ABL1 | 63 |
| TP-008 | ABL1(H396P)-nonphosphorylated | ABL1 | 80 |
| TP-008 | ABL1(H396P)-phosphorylated | ABL1 | 91 |
| TP-008 | ABL1(M351T)-phosphorylated | ABL1 | 81 |
| TP-008 | ABL1(Q252H)-nonphosphorylated | ABL1 | 50 |
| TP-008 | ABL1(Q252H)-phosphorylated | ABL1 | 91 |
| TP-008 | ABL1(T315I)-nonphosphorylated | ABL1 | 77 |
| TP-008 | ABL1(T315I)-phosphorylated | ABL1 | 61 |
| TP-008 | ABL1(Y253F)-phosphorylated | ABL1 | 84 |
| TP-008 | ABL1-nonphosphorylated | ABL1 | 70 |
| TP-008 | ABL1-phosphorylated | ABL1 | 98 |
| TP-008 | ABL2 | ABL2 | 92 |
| TP-008 | ACVR1 | ACVR1 | 87 |
| TP-008 | ACVR1B | ACVR1B | 3,3 |
| TP-008 | ACVR2A | ACVR2A | 100 |
| TP-008 | ACVR2B | ACVR2B | 100 |
| TP-008 | ACVRL1 | ACVRL1 | 80 |
| TP-008 | ADCK3 | CABC1 | 96 |
| TP-008 | ADCK4 | ADCK4 | 69 |
| TP-008 | AKT1 | AKT1 | 85 |
| TP-008 | AKT2 | AKT2 | 100 |
| TP-008 | AKT3 | AKT3 | 100 |
| TP-008 | ALK | ALK | 70 |
| TP-008 | ALK(C1156Y) | ALK | 99 |
| TP-008 | ALK(L1196M) | ALK | 84 |
| TP-008 | AMPK-alpha1 | PRKAA1 | 82 |
| TP-008 | AMPK-alpha2 | PRKAA2 | 95 |
| TP-008 | ANKK1 | ANKK1 | 94 |
| TP-008 | ARK5 | NUAK1 | 55 |
| TP-008 | ASK1 | MAP3K5 | 83 |
| TP-008 | ASK2 | MAP3K6 | 94 |
| TP-008 | AURKA | AURKA | 93 |
| TP-008 | AURKB | AURKB | 84 |
| TP-008 | AURKC | AURKC | 85 |
| TP-008 | AXL | AXL | 100 |
| TP-008 | BIKE | BMP2K | 66 |
| TP-008 | BLK | BLK | 76 |
| TP-008 | BMPR1A | BMPR1A | 98 |
| TP-008 | BMPR1B | BMPR1B | 78 |

| | | | |
|---|---|---|---|
| TP-008 | BMPR2 | BMPR2 | 85 |
| TP-008 | BMX | BMX | 100 |
| TP-008 | BRAF | BRAF | 100 |
| TP-008 | BRAF(V600E) | BRAF | 100 |
| TP-008 | BRK | PTK6 | 100 |
| TP-008 | BRSK1 | BRSK1 | 100 |
| TP-008 | BRSK2 | BRSK2 | 100 |
| TP-008 | BTK | BTK | 77 |
| TP-008 | BUB1 | BUB1 | 100 |
| TP-008 | CAMK1 | CAMK1 | 64 |
| TP-008 | CAMK1B | PNCK | 40 |
| TP-008 | CAMK1D | CAMK1D | 79 |
| TP-008 | CAMK1G | CAMK1G | 100 |
| TP-008 | CAMK2A | CAMK2A | 100 |
| TP-008 | CAMK2B | CAMK2B | 100 |
| TP-008 | CAMK2D | CAMK2D | 90 |
| TP-008 | CAMK2G | CAMK2G | 89 |
| TP-008 | CAMK4 | CAMK4 | 90 |
| TP-008 | CAMKK1 | CAMKK1 | 90 |
| TP-008 | CAMKK2 | CAMKK2 | 88 |
| TP-008 | CASK | CASK | 77 |
| TP-008 | CDC2L1 | CDK11B | 96 |
| TP-008 | CDC2L2 | CDC2L2 | 91 |
| TP-008 | CDC2L5 | CDK13 | 78 |
| TP-008 | CDK11 | CDK19 | 67 |
| TP-008 | CDK2 | CDK2 | 100 |
| TP-008 | CDK3 | CDK3 | 100 |
| TP-008 | CDK4 | CDK4 | 76 |
| TP-008 | CDK4-cyclinD1 | CDK4 | 68 |
| TP-008 | CDK4-cyclinD3 | CDK4 | 100 |
| TP-008 | CDK5 | CDK5 | 79 |
| TP-008 | CDK7 | CDK7 | 83 |
| TP-008 | CDK8 | CDK8 | 97 |
| TP-008 | CDK9 | CDK9 | 91 |
| TP-008 | CDKL1 | CDKL1 | 100 |
| TP-008 | CDKL2 | CDKL2 | 88 |
| TP-008 | CDKL3 | CDKL3 | 78 |
| TP-008 | CDKL5 | CDKL5 | 59 |
| TP-008 | CHEK1 | CHEK1 | 86 |
| TP-008 | CHEK2 | CHEK2 | 100 |
| TP-008 | CIT | CIT | 86 |
| TP-008 | CLK1 | CLK1 | 100 |
| TP-008 | CLK2 | CLK2 | 72 |
| TP-008 | CLK3 | CLK3 | 98 |
| TP-008 | CLK4 | CLK4 | 91 |
| TP-008 | CSF1R | CSF1R | 92 |
| TP-008 | CSF1R-autoinhibited | CSF1R | 66 |

| | | | |
|---|---|---|---|
| TP-008 | CSK | CSK | 80 |
| TP-008 | CSNK1A1 | CSNK1A1 | 71 |
| TP-008 | CSNK1A1L | CSNK1A1L | 97 |
| TP-008 | CSNK1D | CSNK1D | 100 |
| TP-008 | CSNK1E | CSNK1E | 48 |
| TP-008 | CSNK1G1 | CSNK1G1 | 100 |
| TP-008 | CSNK1G2 | CSNK1G2 | 84 |
| TP-008 | CSNK1G3 | CSNK1G3 | 100 |
| TP-008 | CSNK2A1 | CSNK2A1 | 63 |
| TP-008 | CSNK2A2 | CSNK2A2 | 100 |
| TP-008 | CTK | MATK | 100 |
| TP-008 | DAPK1 | DAPK1 | 100 |
| TP-008 | DAPK2 | DAPK2 | 100 |
| TP-008 | DAPK3 | DAPK3 | 96 |
| TP-008 | DCAMKL1 | DCLK1 | 60 |
| TP-008 | DCAMKL2 | DCLK2 | 87 |
| TP-008 | DCAMKL3 | DCLK3 | 82 |
| TP-008 | DDR1 | DDR1 | 37 |
| TP-008 | DDR2 | DDR2 | 92 |
| TP-008 | DLK | MAP3K12 | 52 |
| TP-008 | DMPK | DMPK | 86 |
| TP-008 | DMPK2 | CDC42BPG | 69 |
| TP-008 | DRAK1 | STK17A | 92 |
| TP-008 | DRAK2 | STK17B | 78 |
| TP-008 | DYRK1A | DYRK1A | 63 |
| TP-008 | DYRK1B | DYRK1B | 46 |
| TP-008 | DYRK2 | DYRK2 | 68 |
| TP-008 | EGFR | EGFR | 100 |
| TP-008 | EGFR(E746-A750del) | EGFR | 84 |
| TP-008 | EGFR(G719C) | EGFR | 100 |
| TP-008 | EGFR(G719S) | EGFR | 97 |
| TP-008 | EGFR(L747-E749del, A750P) | EGFR | 77 |
| TP-008 | EGFR(L747-S752del, P753S) | EGFR | 82 |
| TP-008 | EGFR(L747-T751del,Sins) | EGFR | 100 |
| TP-008 | EGFR(L858R) | EGFR | 81 |
| TP-008 | EGFR(L858R,T790M) | EGFR | 95 |
| TP-008 | EGFR(L861Q) | EGFR | 100 |
| TP-008 | EGFR(S752-I759del) | EGFR | 100 |
| TP-008 | EGFR(T790M) | EGFR | 100 |
| TP-008 | EIF2AK1 | EIF2AK1 | 100 |
| TP-008 | EPHA1 | EPHA1 | 100 |
| TP-008 | EPHA2 | EPHA2 | 100 |
| TP-008 | EPHA3 | EPHA3 | 100 |
| TP-008 | EPHA4 | EPHA4 | 83 |
| TP-008 | EPHA5 | EPHA5 | 83 |
| TP-008 | EPHA6 | EPHA6 | 98 |
| TP-008 | EPHA7 | EPHA7 | 84 |

| | | | |
|---|---|---|---|
| TP-008 | EPHA8 | EPHA8 | 100 |
| TP-008 | EPHB1 | EPHB1 | 94 |
| TP-008 | EPHB2 | EPHB2 | 95 |
| TP-008 | EPHB3 | EPHB3 | 100 |
| TP-008 | EPHB4 | EPHB4 | 100 |
| TP-008 | EPHB6 | EPHB6 | 100 |
| TP-008 | ERBB2 | ERBB2 | 100 |
| TP-008 | ERBB3 | ERBB3 | 60 |
| TP-008 | ERBB4 | ERBB4 | 89 |
| TP-008 | ERK1 | MAPK3 | 83 |
| TP-008 | ERK2 | MAPK1 | 100 |
| TP-008 | ERK3 | MAPK6 | 74 |
| TP-008 | ERK4 | MAPK4 | 57 |
| TP-008 | ERK5 | MAPK7 | 74 |
| TP-008 | ERK8 | MAPK15 | 100 |
| TP-008 | ERN1 | ERN1 | 68 |
| TP-008 | FAK | PTK2 | 96 |
| TP-008 | FER | FER | 100 |
| TP-008 | FES | FES | 100 |
| TP-008 | FGFR1 | FGFR1 | 84 |
| TP-008 | FGFR2 | FGFR2 | 92 |
| TP-008 | FGFR3 | FGFR3 | 84 |
| TP-008 | FGFR3(G697C) | FGFR3 | 100 |
| TP-008 | FGFR4 | FGFR4 | 100 |
| TP-008 | FGR | FGR | 100 |
| TP-008 | FLT1 | FLT1 | 93 |
| TP-008 | FLT3 | FLT3 | 100 |
| TP-008 | FLT3(D835H) | FLT3 | 86 |
| TP-008 | FLT3(D835V) | FLT3 | 78 |
| TP-008 | FLT3(D835Y) | FLT3 | 93 |
| TP-008 | FLT3(ITD) | FLT3 | 98 |
| TP-008 | FLT3(ITD,D835V) | FLT3 | 72 |
| TP-008 | FLT3(ITD,F691L) | FLT3 | 93 |
| TP-008 | FLT3(K663Q) | FLT3 | 100 |
| TP-008 | FLT3(N841I) | FLT3 | 68 |
| TP-008 | FLT3(R834Q) | FLT3 | 84 |
| TP-008 | FLT3-autoinhibited | FLT3 | 100 |
| TP-008 | FLT4 | FLT4 | 64 |
| TP-008 | FRK | FRK | 100 |
| TP-008 | FYN | FYN | 100 |
| TP-008 | GAK | GAK | 100 |
| TP-008 | GCN2(Kin.Dom.2,S808G) | EIF2AK4 | 47 |
| TP-008 | GRK1 | GRK1 | 78 |
| TP-008 | GRK2 | ADRBK1 | 64 |
| TP-008 | GRK3 | ADRBK2 | 61 |
| TP-008 | GRK4 | GRK4 | 100 |
| TP-008 | GRK7 | GRK7 | 100 |

| | | | |
|---|---|---|---|
| TP-008 | GSK3A | GSK3A | 84 |
| TP-008 | GSK3B | GSK3B | 68 |
| TP-008 | HASPIN | GSG2 | 81 |
| TP-008 | HCK | HCK | 66 |
| TP-008 | HIPK1 | HIPK1 | 59 |
| TP-008 | HIPK2 | HIPK2 | 78 |
| TP-008 | HIPK3 | HIPK3 | 72 |
| TP-008 | HIPK4 | HIPK4 | 91 |
| TP-008 | HPK1 | MAP4K1 | 89 |
| TP-008 | HUNK | HUNK | 91 |
| TP-008 | ICK | ICK | 79 |
| TP-008 | IGF1R | IGF1R | 100 |
| TP-008 | IKK-alpha | CHUK | 100 |
| TP-008 | IKK-beta | IKBKB | 100 |
| TP-008 | IKK-epsilon | IKBKE | 100 |
| TP-008 | INSR | INSR | 100 |
| TP-008 | INSRR | INSRR | 100 |
| TP-008 | IRAK1 | IRAK1 | 80 |
| TP-008 | IRAK3 | IRAK3 | 83 |
| TP-008 | IRAK4 | IRAK4 | 92 |
| TP-008 | ITK | ITK | 88 |
| TP-008 | JAK1(JH1domain-catalytic) | JAK1 | 98 |
| TP-008 | JAK1(JH2domain-pseudokinase) | JAK1 | 39 |
| TP-008 | JAK2(JH1domain-catalytic) | JAK2 | 85 |
| TP-008 | JAK3(JH1domain-catalytic) | JAK3 | 82 |
| TP-008 | JNK1 | MAPK8 | 58 |
| TP-008 | JNK2 | MAPK9 | 40 |
| TP-008 | JNK3 | MAPK10 | 42 |
| TP-008 | KIT | KIT | 86 |
| TP-008 | KIT(A829P) | KIT | 79 |
| TP-008 | KIT(D816H) | KIT | 64 |
| TP-008 | KIT(D816V) | KIT | 100 |
| TP-008 | KIT(L576P) | KIT | 97 |
| TP-008 | KIT(V559D) | KIT | 86 |
| TP-008 | KIT(V559D,T670I) | KIT | 76 |
| TP-008 | KIT(V559D,V654A) | KIT | 100 |
| TP-008 | KIT-autoinhibited | KIT | 88 |
| TP-008 | LATS1 | LATS1 | 64 |
| TP-008 | LATS2 | LATS2 | 100 |
| TP-008 | LCK | LCK | 88 |
| TP-008 | LIMK1 | LIMK1 | 90 |
| TP-008 | LIMK2 | LIMK2 | 85 |
| TP-008 | LKB1 | STK11 | 60 |
| TP-008 | LOK | STK10 | 66 |
| TP-008 | LRRK2 | LRRK2 | 78 |
| TP-008 | LRRK2(G2019S) | LRRK2 | 73 |
| TP-008 | LTK | LTK | 91 |

| | | | |
|---|---|---|---|
| TP-008 | LYN | LYN | 85 |
| TP-008 | LZK | MAP3K13 | 100 |
| TP-008 | MAK | MAK | 80 |
| TP-008 | MAP3K1 | MAP3K1 | 90 |
| TP-008 | MAP3K15 | MAP3K15 | 81 |
| TP-008 | MAP3K2 | MAP3K2 | 54 |
| TP-008 | MAP3K3 | MAP3K3 | 80 |
| TP-008 | MAP3K4 | MAP3K4 | 71 |
| TP-008 | MAP4K2 | MAP4K2 | 100 |
| TP-008 | MAP4K3 | MAP4K3 | 100 |
| TP-008 | MAP4K4 | MAP4K4 | 82 |
| TP-008 | MAP4K5 | MAP4K5 | 100 |
| TP-008 | MAPKAPK2 | MAPKAPK2 | 100 |
| TP-008 | MAPKAPK5 | MAPKAPK5 | 97 |
| TP-008 | MARK1 | MARK1 | 86 |
| TP-008 | MARK2 | MARK2 | 93 |
| TP-008 | MARK3 | MARK3 | 100 |
| TP-008 | MARK4 | MARK4 | 97 |
| TP-008 | MAST1 | MAST1 | 100 |
| TP-008 | MEK1 | MAP2K1 | 100 |
| TP-008 | MEK2 | MAP2K2 | 97 |
| TP-008 | MEK3 | MAP2K3 | 61 |
| TP-008 | MEK4 | MAP2K4 | 95 |
| TP-008 | MEK5 | MAP2K5 | 53 |
| TP-008 | MEK6 | MAP2K6 | 87 |
| TP-008 | MELK | MELK | 80 |
| TP-008 | MERTK | MERTK | 78 |
| TP-008 | MET | MET | 68 |
| TP-008 | MET(M1250T) | MET | 68 |
| TP-008 | MET(Y1235D) | MET | 97 |
| TP-008 | MINK | MINK1 | 61 |
| TP-008 | MKK7 | MAP2K7 | 73 |
| TP-008 | MKNK1 | MKNK1 | 76 |
| TP-008 | MKNK2 | MKNK2 | 74 |
| TP-008 | MLCK | MYLK3 | 100 |
| TP-008 | MLK1 | MAP3K9 | 81 |
| TP-008 | MLK2 | MAP3K10 | 100 |
| TP-008 | MLK3 | MAP3K11 | 96 |
| TP-008 | MRCKA | CDC42BPA | 76 |
| TP-008 | MRCKB | CDC42BPB | 100 |
| TP-008 | MST1 | STK4 | 100 |
| TP-008 | MST1R | MST1R | 63 |
| TP-008 | MST2 | STK3 | 85 |
| TP-008 | MST3 | STK24 | 89 |
| TP-008 | MST4 | MST4 | 100 |
| TP-008 | MTOR | MTOR | 82 |
| TP-008 | MUSK | MUSK | 100 |

| | | | |
|---|---|---|---|
| TP-008 | MYLK | MYLK | 83 |
| TP-008 | MYLK2 | MYLK2 | 93 |
| TP-008 | MYLK4 | MYLK4 | 91 |
| TP-008 | MYO3A | MYO3A | 100 |
| TP-008 | MYO3B | MYO3B | 69 |
| TP-008 | NDR1 | STK38 | 96 |
| TP-008 | NDR2 | STK38L | 100 |
| TP-008 | NEK1 | NEK1 | 85 |
| TP-008 | NEK10 | NEK10 | 45 |
| TP-008 | NEK11 | NEK11 | 49 |
| TP-008 | NEK2 | NEK2 | 80 |
| TP-008 | NEK3 | NEK3 | 62 |
| TP-008 | NEK4 | NEK4 | 91 |
| TP-008 | NEK5 | NEK5 | 68 |
| TP-008 | NEK6 | NEK6 | 73 |
| TP-008 | NEK7 | NEK7 | 90 |
| TP-008 | NEK9 | NEK9 | 93 |
| TP-008 | NIK | MAP3K14 | 92 |
| TP-008 | NIM1 | MGC42105 | 100 |
| TP-008 | NLK | NLK | 100 |
| TP-008 | OSR1 | OXSR1 | 98 |
| TP-008 | p38-alpha | MAPK14 | 97 |
| TP-008 | p38-beta | MAPK11 | 86 |
| TP-008 | p38-delta | MAPK13 | 100 |
| TP-008 | p38-gamma | MAPK12 | 60 |
| TP-008 | PAK1 | PAK1 | 91 |
| TP-008 | PAK2 | PAK2 | 91 |
| TP-008 | PAK3 | PAK3 | 73 |
| TP-008 | PAK4 | PAK4 | 85 |
| TP-008 | PAK6 | PAK6 | 100 |
| TP-008 | PAK7 | PAK7 | 72 |
| TP-008 | PCTK1 | CDK16 | 78 |
| TP-008 | PCTK2 | CDK17 | 88 |
| TP-008 | PCTK3 | CDK18 | 74 |
| TP-008 | PDGFRA | PDGFRA | 100 |
| TP-008 | PDGFRB | PDGFRB | 65 |
| TP-008 | PDPK1 | PDPK1 | 84 |
| TP-008 | PFCDPK1(P.falciparum) | CDPK1 | 70 |
| TP-008 | PFPK5(P.falciparum) | MAL13P1.279 | 84 |
| TP-008 | PFTAIRE2 | CDK15 | 65 |
| TP-008 | PFTK1 | CDK14 | 100 |
| TP-008 | PHKG1 | PHKG1 | 92 |
| TP-008 | PHKG2 | PHKG2 | 100 |
| TP-008 | PIK3C2B | PIK3C2B | 100 |
| TP-008 | PIK3C2G | PIK3C2G | 78 |
| TP-008 | PIK3CA | PIK3CA | 100 |
| TP-008 | PIK3CA(C420R) | PIK3CA | 100 |

| | | | |
|---|---|---|---|
| TP-008 | PIK3CA(E542K) | PIK3CA | 76 |
| TP-008 | PIK3CA(E545A) | PIK3CA | 100 |
| TP-008 | PIK3CA(E545K) | PIK3CA | 85 |
| TP-008 | PIK3CA(H1047L) | PIK3CA | 98 |
| TP-008 | PIK3CA(H1047Y) | PIK3CA | 58 |
| TP-008 | PIK3CA(I800L) | PIK3CA | 100 |
| TP-008 | PIK3CA(M1043I) | PIK3CA | 34 |
| TP-008 | PIK3CA(Q546K) | PIK3CA | 100 |
| TP-008 | PIK3CB | PIK3CB | 100 |
| TP-008 | PIK3CD | PIK3CD | 100 |
| TP-008 | PIK3CG | PIK3CG | 100 |
| TP-008 | PIK4CB | PI4KB | 100 |
| TP-008 | PIKFYVE | PIKFYVE | 97 |
| TP-008 | PIM1 | PIM1 | 94 |
| TP-008 | PIM2 | PIM2 | 81 |
| TP-008 | PIM3 | PIM3 | 100 |
| TP-008 | PIP5K1A | PIP5K1A | 100 |
| TP-008 | PIP5K1C | PIP5K1C | 100 |
| TP-008 | PIP5K2B | PIP4K2B | 60 |
| TP-008 | PIP5K2C | PIP4K2C | 100 |
| TP-008 | PKAC-alpha | PRKACA | 73 |
| TP-008 | PKAC-beta | PRKACB | 100 |
| TP-008 | PKMYT1 | PKMYT1 | 100 |
| TP-008 | PKN1 | PKN1 | 89 |
| TP-008 | PKN2 | PKN2 | 100 |
| TP-008 | PKNB(M.tuberculosis) | pknB | 100 |
| TP-008 | PLK1 | PLK1 | 64 |
| TP-008 | PLK2 | PLK2 | 100 |
| TP-008 | PLK3 | PLK3 | 73 |
| TP-008 | PLK4 | PLK4 | 100 |
| TP-008 | PRKCD | PRKCD | 77 |
| TP-008 | PRKCE | PRKCE | 100 |
| TP-008 | PRKCH | PRKCH | 89 |
| TP-008 | PRKCI | PRKCI | 100 |
| TP-008 | PRKCQ | PRKCQ | 62 |
| TP-008 | PRKD1 | PRKD1 | 97 |
| TP-008 | PRKD2 | PRKD2 | 100 |
| TP-008 | PRKD3 | PRKD3 | 80 |
| TP-008 | PRKG1 | PRKG1 | 100 |
| TP-008 | PRKG2 | PRKG2 | 100 |
| TP-008 | PRKR | EIF2AK2 | 69 |
| TP-008 | PRKX | PRKX | 74 |
| TP-008 | PRP4 | PRPF4B | 100 |
| TP-008 | PYK2 | PTK2B | 100 |
| TP-008 | QSK | KIAA0999 | 79 |
| TP-008 | RAF1 | RAF1 | 100 |
| TP-008 | RET | RET | 65 |

| | | | |
|---|---|---|---|
| TP-008 | RET(M918T) | RET | 88 |
| TP-008 | RET(V804L) | RET | 90 |
| TP-008 | RET(V804M) | RET | 80 |
| TP-008 | RIOK1 | RIOK1 | 93 |
| TP-008 | RIOK2 | RIOK2 | 94 |
| TP-008 | RIOK3 | RIOK3 | 84 |
| TP-008 | RIPK1 | RIPK1 | 67 |
| TP-008 | RIPK2 | RIPK2 | 93 |
| TP-008 | RIPK4 | RIPK4 | 48 |
| TP-008 | RIPK5 | DSTYK | 73 |
| TP-008 | ROCK1 | ROCK1 | 59 |
| TP-008 | ROCK2 | ROCK2 | 65 |
| TP-008 | ROS1 | ROS1 | 100 |
| TP-008 | RPS6KA4(Kin.Dom.1-N-terminal) | RPS6KA4 | 100 |
| TP-008 | RPS6KA4(Kin.Dom.2-C-terminal) | RPS6KA4 | 100 |
| TP-008 | RPS6KA5(Kin.Dom.1-N-terminal) | RPS6KA5 | 100 |
| TP-008 | RPS6KA5(Kin.Dom.2-C-terminal) | RPS6KA5 | 100 |
| TP-008 | RSK1(Kin.Dom.1-N-terminal) | RPS6KA1 | 86 |
| TP-008 | RSK1(Kin.Dom.2-C-terminal) | RPS6KA1 | 100 |
| TP-008 | RSK2(Kin.Dom.1-N-terminal) | RPS6KA3 | 84 |
| TP-008 | RSK2(Kin.Dom.2-C-terminal) | RPS6KA3 | 74 |
| TP-008 | RSK3(Kin.Dom.1-N-terminal) | RPS6KA2 | 90 |
| TP-008 | RSK3(Kin.Dom.2-C-terminal) | RPS6KA2 | 93 |
| TP-008 | RSK4(Kin.Dom.1-N-terminal) | RPS6KA6 | 100 |
| TP-008 | RSK4(Kin.Dom.2-C-terminal) | RPS6KA6 | 92 |
| TP-008 | S6K1 | RPS6KB1 | 80 |
| TP-008 | SBK1 | SBK1 | 100 |
| TP-008 | SGK | SGK1 | 74 |
| TP-008 | SgK110 | SgK110 | 86 |
| TP-008 | SGK2 | SGK2 | 81 |
| TP-008 | SGK3 | SGK3 | 65 |
| TP-008 | SIK | SIK1 | 78 |
| TP-008 | SIK2 | SIK2 | 99 |
| TP-008 | SLK | SLK | 96 |
| TP-008 | SNARK | NUAK2 | 100 |
| TP-008 | SNRK | SNRK | 90 |
| TP-008 | SRC | SRC | 91 |
| TP-008 | SRMS | SRMS | 100 |
| TP-008 | SRPK1 | SRPK1 | 99 |
| TP-008 | SRPK2 | SRPK2 | 100 |
| TP-008 | SRPK3 | SRPK3 | 100 |
| TP-008 | STK16 | STK16 | 75 |
| TP-008 | STK33 | STK33 | 56 |
| TP-008 | STK35 | STK35 | 100 |
| TP-008 | STK36 | STK36 | 68 |
| TP-008 | STK39 | STK39 | 88 |
| TP-008 | SYK | SYK | 82 |

| | | | |
|---|---|---|---|
| TP-008 | TAK1 | MAP3K7 | 100 |
| TP-008 | TAOK1 | TAOK1 | 100 |
| TP-008 | TAOK2 | TAOK2 | 100 |
| TP-008 | TAOK3 | TAOK3 | 100 |
| TP-008 | TBK1 | TBK1 | 82 |
| TP-008 | TEC | TEC | 90 |
| TP-008 | TESK1 | TESK1 | 96 |
| TP-008 | TGFBR1 | TGFBR1 | 3,6 |
| TP-008 | TGFBR2 | TGFBR2 | 83 |
| TP-008 | TIE1 | TIE1 | 100 |
| TP-008 | TIE2 | TEK | 77 |
| TP-008 | TLK1 | TLK1 | 66 |
| TP-008 | TLK2 | TLK2 | 96 |
| TP-008 | TNIK | TNIK | 97 |
| TP-008 | TNK1 | TNK1 | 58 |
| TP-008 | TNK2 | TNK2 | 100 |
| TP-008 | TNNI3K | TNNI3K | 100 |
| TP-008 | TRKA | NTRK1 | 70 |
| TP-008 | TRKB | NTRK2 | 55 |
| TP-008 | TRKC | NTRK3 | 81 |
| TP-008 | TRPM6 | TRPM6 | 92 |
| TP-008 | TSSK1B | TSSK1B | 83 |
| TP-008 | TSSK3 | TSSK3 | 100 |
| TP-008 | TTK | TTK | 67 |
| TP-008 | TXK | TXK | 59 |
| TP-008 | TYK2(JH1domain-catalytic) | TYK2 | 66 |
| TP-008 | TYK2(JH2domain-pseudokinase) | TYK2 | 100 |
| TP-008 | TYRO3 | TYRO3 | 86 |
| TP-008 | ULK1 | ULK1 | 100 |
| TP-008 | ULK2 | ULK2 | 90 |
| TP-008 | ULK3 | ULK3 | 94 |
| TP-008 | VEGFR2 | KDR | 89 |
| TP-008 | VPS34 | PIK3C3 | 68 |
| TP-008 | VRK2 | VRK2 | 59 |
| TP-008 | WEE1 | WEE1 | 96 |
| TP-008 | WEE2 | WEE2 | 93 |
| TP-008 | WNK1 | WNK1 | 100 |
| TP-008 | WNK2 | WNK2 | 51 |
| TP-008 | WNK3 | WNK3 | 100 |
| TP-008 | WNK4 | WNK4 | 100 |
| TP-008 | YANK1 | STK32A | 100 |
| TP-008 | YANK2 | STK32B | 88 |
| TP-008 | YANK3 | STK32C | 100 |
| TP-008 | YES | YES1 | 100 |
| TP-008 | YSK1 | STK25 | 76 |
| TP-008 | YSK4 | MAP3K19 | 92 |
| TP-008 | ZAK | ZAK | 85 |

| | | | |
|---|---|---|---|
| TP-008 | ZAP70 | ZAP70 | 88 |

**Table S6**. Kinome scan of THAI11 (**15**) @ 1 µM against 469 kinases from DiscoverX/Eurofins.

| Compound Name | DiscoveRx Gene Symbol | Entrez Gene Symbol | Percent Control |
|---|---|---|---|
| THAI11 | AAK1 | AAK1 | 62 |
| THAI11 | ABL1(E255K)-phosphorylated | ABL1 | 93 |
| THAI11 | ABL1(F317I)-nonphosphorylated | ABL1 | 82 |
| THAI11 | ABL1(F317I)-phosphorylated | ABL1 | 93 |
| THAI11 | ABL1(F317L)-nonphosphorylated | ABL1 | 74 |
| THAI11 | ABL1(F317L)-phosphorylated | ABL1 | 75 |
| THAI11 | ABL1(H396P)-nonphosphorylated | ABL1 | 78 |
| THAI11 | ABL1(H396P)-phosphorylated | ABL1 | 83 |
| THAI11 | ABL1(M351T)-phosphorylated | ABL1 | 73 |
| THAI11 | ABL1(Q252H)-nonphosphorylated | ABL1 | 89 |
| THAI11 | ABL1(Q252H)-phosphorylated | ABL1 | 100 |
| THAI11 | ABL1(T315I)-nonphosphorylated | ABL1 | 81 |
| THAI11 | ABL1(T315I)-phosphorylated | ABL1 | 77 |
| THAI11 | ABL1(Y253F)-phosphorylated | ABL1 | 88 |
| THAI11 | ABL1-nonphosphorylated | ABL1 | 58 |
| THAI11 | ABL1-phosphorylated | ABL1 | 77 |
| THAI11 | ABL2 | ABL2 | 99 |
| THAI11 | ACVR1 | ACVR1 | 88 |
| THAI11 | ACVR1B | ACVR1B | 88 |
| THAI11 | ACVR2A | ACVR2A | 83 |
| THAI11 | ACVR2B | ACVR2B | 91 |
| THAI11 | ACVRL1 | ACVRL1 | 76 |
| THAI11 | ADCK3 | CABC1 | 82 |
| THAI11 | ADCK4 | ADCK4 | 74 |
| THAI11 | AKT1 | AKT1 | 78 |
| THAI11 | AKT2 | AKT2 | 86 |
| THAI11 | AKT3 | AKT3 | 91 |
| THAI11 | ALK | ALK | 70 |
| THAI11 | ALK(C1156Y) | ALK | 94 |
| THAI11 | ALK(L1196M) | ALK | 62 |
| THAI11 | AMPK-alpha1 | PRKAA1 | 87 |
| THAI11 | AMPK-alpha2 | PRKAA2 | 100 |
| THAI11 | ANKK1 | ANKK1 | 85 |
| THAI11 | ARK5 | NUAK1 | 85 |
| THAI11 | ASK1 | MAP3K5 | 89 |
| THAI11 | ASK2 | MAP3K6 | 94 |
| THAI11 | AURKA | AURKA | 98 |

| | | | |
|---|---|---|---|
| THAI11 | AURKB | AURKB | 97 |
| THAI11 | AURKC | AURKC | 82 |
| THAI11 | AXL | AXL | 100 |
| THAI11 | BIKE | BMP2K | 83 |
| THAI11 | BLK | BLK | 99 |
| THAI11 | BMPR1A | BMPR1A | 88 |
| THAI11 | BMPR1B | BMPR1B | 76 |
| THAI11 | BMPR2 | BMPR2 | 95 |
| THAI11 | BMX | BMX | 83 |
| THAI11 | BRAF | BRAF | 86 |
| THAI11 | BRAF(V600E) | BRAF | 79 |
| THAI11 | BRK | PTK6 | 72 |
| THAI11 | BRSK1 | BRSK1 | 100 |
| THAI11 | BRSK2 | BRSK2 | 100 |
| THAI11 | BTK | BTK | 58 |
| THAI11 | BUB1 | BUB1 | 62 |
| THAI11 | CAMK1 | CAMK1 | 67 |
| THAI11 | CAMK1B | PNCK | 73 |
| THAI11 | CAMK1D | CAMK1D | 78 |
| THAI11 | CAMK1G | CAMK1G | 81 |
| THAI11 | CAMK2A | CAMK2A | 71 |
| THAI11 | CAMK2B | CAMK2B | 83 |
| THAI11 | CAMK2D | CAMK2D | 79 |
| THAI11 | CAMK2G | CAMK2G | 93 |
| THAI11 | CAMK4 | CAMK4 | 81 |
| THAI11 | CAMKK1 | CAMKK1 | 93 |
| THAI11 | CAMKK2 | CAMKK2 | 97 |
| THAI11 | CASK | CASK | 89 |
| THAI11 | CDC2L1 | CDK11B | 82 |
| THAI11 | CDC2L2 | CDC2L2 | 100 |
| THAI11 | CDC2L5 | CDK13 | 100 |
| THAI11 | CDK11 | CDK19 | 100 |
| THAI11 | CDK2 | CDK2 | 100 |
| THAI11 | CDK3 | CDK3 | 86 |
| THAI11 | CDK4 | CDK4 | 100 |
| THAI11 | CDK4-cyclinD1 | CDK4 | 78 |
| THAI11 | CDK4-cyclinD3 | CDK4 | 96 |
| THAI11 | CDK5 | CDK5 | 93 |
| THAI11 | CDK7 | CDK7 | 76 |
| THAI11 | CDK8 | CDK8 | 100 |
| THAI11 | CDK9 | CDK9 | 85 |
| THAI11 | CDKL1 | CDKL1 | 83 |
| THAI11 | CDKL2 | CDKL2 | 100 |
| THAI11 | CDKL3 | CDKL3 | 92 |

| | | | |
|---|---|---|---|
| THAI11 | CDKL5 | CDKL5 | 76 |
| THAI11 | CHEK1 | CHEK1 | 77 |
| THAI11 | CHEK2 | CHEK2 | 100 |
| THAI11 | CIT | CIT | 78 |
| THAI11 | CLK1 | CLK1 | 82 |
| THAI11 | CLK2 | CLK2 | 89 |
| THAI11 | CLK3 | CLK3 | 88 |
| THAI11 | CLK4 | CLK4 | 100 |
| THAI11 | CSF1R | CSF1R | 82 |
| THAI11 | CSF1R-autoinhibited | CSF1R | 70 |
| THAI11 | CSK | CSK | 81 |
| THAI11 | CSNK1A1 | CSNK1A1 | 99 |
| THAI11 | CSNK1A1L | CSNK1A1L | 89 |
| THAI11 | CSNK1D | CSNK1D | 100 |
| THAI11 | CSNK1E | CSNK1E | 100 |
| THAI11 | CSNK1G1 | CSNK1G1 | 97 |
| THAI11 | CSNK1G2 | CSNK1G2 | 76 |
| THAI11 | CSNK1G3 | CSNK1G3 | 100 |
| THAI11 | CSNK2A1 | CSNK2A1 | 76 |
| THAI11 | CSNK2A2 | CSNK2A2 | 100 |
| THAI11 | CTK | MATK | 100 |
| THAI11 | DAPK1 | DAPK1 | 89 |
| THAI11 | DAPK2 | DAPK2 | 87 |
| THAI11 | DAPK3 | DAPK3 | 78 |
| THAI11 | DCAMKL1 | DCLK1 | 68 |
| THAI11 | DCAMKL2 | DCLK2 | 95 |
| THAI11 | DCAMKL3 | DCLK3 | 100 |
| THAI11 | DDR1 | DDR1 | 85 |
| THAI11 | DDR2 | DDR2 | 94 |
| THAI11 | DLK | MAP3K12 | 92 |
| THAI11 | DMPK | DMPK | 96 |
| THAI11 | DMPK2 | CDC42BPG | 81 |
| THAI11 | DRAK1 | STK17A | 100 |
| THAI11 | DRAK2 | STK17B | 91 |
| THAI11 | DYRK1A | DYRK1A | 80 |
| THAI11 | DYRK1B | DYRK1B | 100 |
| THAI11 | DYRK2 | DYRK2 | 69 |
| THAI11 | EGFR | EGFR | 81 |
| THAI11 | EGFR(E746-A750del) | EGFR | 83 |
| THAI11 | EGFR(G719C) | EGFR | 78 |
| THAI11 | EGFR(G719S) | EGFR | 71 |
| THAI11 | EGFR(L747-E749del, A750P) | EGFR | 98 |
| THAI11 | EGFR(L747-S752del, P753S) | EGFR | 81 |
| THAI11 | EGFR(L747-T751del,Sins) | EGFR | 67 |

| | | | |
|---|---|---|---|
| THAI11 | EGFR(L858R) | EGFR | 87 |
| THAI11 | EGFR(L858R,T790M) | EGFR | 86 |
| THAI11 | EGFR(L861Q) | EGFR | 56 |
| THAI11 | EGFR(S752-I759del) | EGFR | 57 |
| THAI11 | EGFR(T790M) | EGFR | 71 |
| THAI11 | EIF2AK1 | EIF2AK1 | 100 |
| THAI11 | EPHA1 | EPHA1 | 99 |
| THAI11 | EPHA2 | EPHA2 | 95 |
| THAI11 | EPHA3 | EPHA3 | 86 |
| THAI11 | EPHA4 | EPHA4 | 94 |
| THAI11 | EPHA5 | EPHA5 | 91 |
| THAI11 | EPHA6 | EPHA6 | 69 |
| THAI11 | EPHA7 | EPHA7 | 100 |
| THAI11 | EPHA8 | EPHA8 | 100 |
| THAI11 | EPHB1 | EPHB1 | 100 |
| THAI11 | EPHB2 | EPHB2 | 75 |
| THAI11 | EPHB3 | EPHB3 | 70 |
| THAI11 | EPHB4 | EPHB4 | 75 |
| THAI11 | EPHB6 | EPHB6 | 84 |
| THAI11 | ERBB2 | ERBB2 | 89 |
| THAI11 | ERBB3 | ERBB3 | 80 |
| THAI11 | ERBB4 | ERBB4 | 77 |
| THAI11 | ERK1 | MAPK3 | 71 |
| THAI11 | ERK2 | MAPK1 | 82 |
| THAI11 | ERK3 | MAPK6 | 80 |
| THAI11 | ERK4 | MAPK4 | 83 |
| THAI11 | ERK5 | MAPK7 | 97 |
| THAI11 | ERK8 | MAPK15 | 98 |
| THAI11 | ERN1 | ERN1 | 67 |
| THAI11 | FAK | PTK2 | 99 |
| THAI11 | FER | FER | 65 |
| THAI11 | FES | FES | 63 |
| THAI11 | FGFR1 | FGFR1 | 100 |
| THAI11 | FGFR2 | FGFR2 | 100 |
| THAI11 | FGFR3 | FGFR3 | 90 |
| THAI11 | FGFR3(G697C) | FGFR3 | 95 |
| THAI11 | FGFR4 | FGFR4 | 75 |
| THAI11 | FGR | FGR | 63 |
| THAI11 | FLT1 | FLT1 | 79 |
| THAI11 | FLT3 | FLT3 | 99 |
| THAI11 | FLT3(D835H) | FLT3 | 90 |
| THAI11 | FLT3(D835V) | FLT3 | 90 |
| THAI11 | FLT3(D835Y) | FLT3 | 83 |
| THAI11 | FLT3(ITD) | FLT3 | 87 |

| | | | |
|---|---|---|---|
| THAI11 | FLT3(ITD,D835V) | FLT3 | 64 |
| THAI11 | FLT3(ITD,F691L) | FLT3 | 80 |
| THAI11 | FLT3(K663Q) | FLT3 | 96 |
| THAI11 | FLT3(N841I) | FLT3 | 97 |
| THAI11 | FLT3(R834Q) | FLT3 | 100 |
| THAI11 | FLT3-autoinhibited | FLT3 | 93 |
| THAI11 | FLT4 | FLT4 | 81 |
| THAI11 | FRK | FRK | 76 |
| THAI11 | FYN | FYN | 77 |
| THAI11 | GAK | GAK | 88 |
| THAI11 | GCN2(Kin.Dom.2,S808G) | EIF2AK4 | 100 |
| THAI11 | GRK1 | GRK1 | 100 |
| THAI11 | GRK2 | ADRBK1 | 68 |
| THAI11 | GRK3 | ADRBK2 | 78 |
| THAI11 | GRK4 | GRK4 | 76 |
| THAI11 | GRK7 | GRK7 | 85 |
| THAI11 | GSK3A | GSK3A | 94 |
| THAI11 | GSK3B | GSK3B | 75 |
| THAI11 | HASPIN | GSG2 | 81 |
| THAI11 | HCK | HCK | 93 |
| THAI11 | HIPK1 | HIPK1 | 71 |
| THAI11 | HIPK2 | HIPK2 | 100 |
| THAI11 | HIPK3 | HIPK3 | 80 |
| THAI11 | HIPK4 | HIPK4 | 73 |
| THAI11 | HPK1 | MAP4K1 | 85 |
| THAI11 | HUNK | HUNK | 82 |
| THAI11 | ICK | ICK | 100 |
| THAI11 | IGF1R | IGF1R | 86 |
| THAI11 | IKK-alpha | CHUK | 82 |
| THAI11 | IKK-beta | IKBKB | 75 |
| THAI11 | IKK-epsilon | IKBKE | 91 |
| THAI11 | INSR | INSR | 69 |
| THAI11 | INSRR | INSRR | 100 |
| THAI11 | IRAK1 | IRAK1 | 93 |
| THAI11 | IRAK3 | IRAK3 | 92 |
| THAI11 | IRAK4 | IRAK4 | 77 |
| THAI11 | ITK | ITK | 100 |
| THAI11 | JAK1(JH1domain-catalytic) | JAK1 | 94 |
| THAI11 | JAK1(JH2domain-pseudokinase) | JAK1 | 100 |
| THAI11 | JAK2(JH1domain-catalytic) | JAK2 | 69 |
| THAI11 | JAK3(JH1domain-catalytic) | JAK3 | 79 |
| THAI11 | JNK1 | MAPK8 | 75 |
| THAI11 | JNK2 | MAPK9 | 66 |
| THAI11 | JNK3 | MAPK10 | 75 |

| | | | |
|---|---|---|---|
| THAI11 | KIT | KIT | 72 |
| THAI11 | KIT(A829P) | KIT | 71 |
| THAI11 | KIT(D816H) | KIT | 73 |
| THAI11 | KIT(D816V) | KIT | 90 |
| THAI11 | KIT(L576P) | KIT | 78 |
| THAI11 | KIT(V559D) | KIT | 88 |
| THAI11 | KIT(V559D,T670I) | KIT | 87 |
| THAI11 | KIT(V559D,V654A) | KIT | 80 |
| THAI11 | KIT-autoinhibited | KIT | 79 |
| THAI11 | LATS1 | LATS1 | 87 |
| THAI11 | LATS2 | LATS2 | 85 |
| THAI11 | LCK | LCK | 79 |
| THAI11 | LIMK1 | LIMK1 | 96 |
| THAI11 | LIMK2 | LIMK2 | 91 |
| THAI11 | LKB1 | STK11 | 100 |
| THAI11 | LOK | STK10 | 84 |
| THAI11 | LRRK2 | LRRK2 | 100 |
| THAI11 | LRRK2(G2019S) | LRRK2 | 100 |
| THAI11 | LTK | LTK | 88 |
| THAI11 | LYN | LYN | 92 |
| THAI11 | LZK | MAP3K13 | 86 |
| THAI11 | MAK | MAK | 87 |
| THAI11 | MAP3K1 | MAP3K1 | 67 |
| THAI11 | MAP3K15 | MAP3K15 | 79 |
| THAI11 | MAP3K2 | MAP3K2 | 82 |
| THAI11 | MAP3K3 | MAP3K3 | 76 |
| THAI11 | MAP3K4 | MAP3K4 | 100 |
| THAI11 | MAP4K2 | MAP4K2 | 100 |
| THAI11 | MAP4K3 | MAP4K3 | 100 |
| THAI11 | MAP4K4 | MAP4K4 | 100 |
| THAI11 | MAP4K5 | MAP4K5 | 100 |
| THAI11 | MAPKAPK2 | MAPKAPK2 | 79 |
| THAI11 | MAPKAPK5 | MAPKAPK5 | 94 |
| THAI11 | MARK1 | MARK1 | 86 |
| THAI11 | MARK2 | MARK2 | 86 |
| THAI11 | MARK3 | MARK3 | 100 |
| THAI11 | MARK4 | MARK4 | 79 |
| THAI11 | MAST1 | MAST1 | 100 |
| THAI11 | MEK1 | MAP2K1 | 94 |
| THAI11 | MEK2 | MAP2K2 | 91 |
| THAI11 | MEK3 | MAP2K3 | 83 |
| THAI11 | MEK4 | MAP2K4 | 88 |
| THAI11 | MEK5 | MAP2K5 | 95 |
| THAI11 | MEK6 | MAP2K6 | 86 |

| | | | |
|---|---|---|---|
| THAI11 | MELK | MELK | 100 |
| THAI11 | MERTK | MERTK | 85 |
| THAI11 | MET | MET | 79 |
| THAI11 | MET(M1250T) | MET | 100 |
| THAI11 | MET(Y1235D) | MET | 100 |
| THAI11 | MINK | MINK1 | 81 |
| THAI11 | MKK7 | MAP2K7 | 89 |
| THAI11 | MKNK1 | MKNK1 | 78 |
| THAI11 | MKNK2 | MKNK2 | 76 |
| THAI11 | MLCK | MYLK3 | 83 |
| THAI11 | MLK1 | MAP3K9 | 76 |
| THAI11 | MLK2 | MAP3K10 | 99 |
| THAI11 | MLK3 | MAP3K11 | 77 |
| THAI11 | MRCKA | CDC42BPA | 88 |
| THAI11 | MRCKB | CDC42BPB | 93 |
| THAI11 | MST1 | STK4 | 81 |
| THAI11 | MST1R | MST1R | 100 |
| THAI11 | MST2 | STK3 | 70 |
| THAI11 | MST3 | STK24 | 100 |
| THAI11 | MST4 | MST4 | 83 |
| THAI11 | MTOR | MTOR | 75 |
| THAI11 | MUSK | MUSK | 100 |
| THAI11 | MYLK | MYLK | 74 |
| THAI11 | MYLK2 | MYLK2 | 90 |
| THAI11 | MYLK4 | MYLK4 | 97 |
| THAI11 | MYO3A | MYO3A | 79 |
| THAI11 | MYO3B | MYO3B | 49 |
| THAI11 | NDR1 | STK38 | 83 |
| THAI11 | NDR2 | STK38L | 68 |
| THAI11 | NEK1 | NEK1 | 95 |
| THAI11 | NEK10 | NEK10 | 85 |
| THAI11 | NEK11 | NEK11 | 100 |
| THAI11 | NEK2 | NEK2 | 100 |
| THAI11 | NEK3 | NEK3 | 87 |
| THAI11 | NEK4 | NEK4 | 97 |
| THAI11 | NEK5 | NEK5 | 90 |
| THAI11 | NEK6 | NEK6 | 85 |
| THAI11 | NEK7 | NEK7 | 72 |
| THAI11 | NEK9 | NEK9 | 89 |
| THAI11 | NIK | MAP3K14 | 82 |
| THAI11 | NIM1 | MGC42105 | 100 |
| THAI11 | NLK | NLK | 100 |
| THAI11 | OSR1 | OXSR1 | 84 |
| THAI11 | p38-alpha | MAPK14 | 100 |

| | | | |
|---|---|---|---|
| THAI11 | p38-beta | MAPK11 | 93 |
| THAI11 | p38-delta | MAPK13 | 91 |
| THAI11 | p38-gamma | MAPK12 | 59 |
| THAI11 | PAK1 | PAK1 | 87 |
| THAI11 | PAK2 | PAK2 | 65 |
| THAI11 | PAK3 | PAK3 | 87 |
| THAI11 | PAK4 | PAK4 | 97 |
| THAI11 | PAK6 | PAK6 | 100 |
| THAI11 | PAK7 | PAK7 | 100 |
| THAI11 | PCTK1 | CDK16 | 83 |
| THAI11 | PCTK2 | CDK17 | 77 |
| THAI11 | PCTK3 | CDK18 | 88 |
| THAI11 | PDGFRA | PDGFRA | 99 |
| THAI11 | PDGFRB | PDGFRB | 69 |
| THAI11 | PDPK1 | PDPK1 | 86 |
| THAI11 | PFCDPK1(P.falciparum) | CDPK1 | 83 |
| THAI11 | PFPK5(P.falciparum) | MAL13P1.279 | 80 |
| THAI11 | PFTAIRE2 | CDK15 | 84 |
| THAI11 | PFTK1 | CDK14 | 100 |
| THAI11 | PHKG1 | PHKG1 | 96 |
| THAI11 | PHKG2 | PHKG2 | 87 |
| THAI11 | PIK3C2B | PIK3C2B | 100 |
| THAI11 | PIK3C2G | PIK3C2G | 100 |
| THAI11 | PIK3CA | PIK3CA | 100 |
| THAI11 | PIK3CA(C420R) | PIK3CA | 97 |
| THAI11 | PIK3CA(E542K) | PIK3CA | 47 |
| THAI11 | PIK3CA(E545A) | PIK3CA | 100 |
| THAI11 | PIK3CA(E545K) | PIK3CA | 100 |
| THAI11 | PIK3CA(H1047L) | PIK3CA | 100 |
| THAI11 | PIK3CA(H1047Y) | PIK3CA | 100 |
| THAI11 | PIK3CA(I800L) | PIK3CA | 100 |
| THAI11 | PIK3CA(M1043I) | PIK3CA | 100 |
| THAI11 | PIK3CA(Q546K) | PIK3CA | 41 |
| THAI11 | PIK3CB | PIK3CB | 100 |
| THAI11 | PIK3CD | PIK3CD | 100 |
| THAI11 | PIK3CG | PIK3CG | 100 |
| THAI11 | PIK4CB | PI4KB | 100 |
| THAI11 | PIKFYVE | PIKFYVE | 73 |
| THAI11 | PIM1 | PIM1 | 95 |
| THAI11 | PIM2 | PIM2 | 84 |
| THAI11 | PIM3 | PIM3 | 94 |
| THAI11 | PIP5K1A | PIP5K1A | 100 |
| THAI11 | PIP5K1C | PIP5K1C | 80 |
| THAI11 | PIP5K2B | PIP4K2B | 96 |

| | | | |
|---|---|---|---|
| THAI11 | PIP5K2C | PIP4K2C | 100 |
| THAI11 | PKAC-alpha | PRKACA | 93 |
| THAI11 | PKAC-beta | PRKACB | 93 |
| THAI11 | PKMYT1 | PKMYT1 | 98 |
| THAI11 | PKN1 | PKN1 | 86 |
| THAI11 | PKN2 | PKN2 | 100 |
| THAI11 | PKNB(M.tuberculosis) | pknB | 80 |
| THAI11 | PLK1 | PLK1 | 86 |
| THAI11 | PLK2 | PLK2 | 82 |
| THAI11 | PLK3 | PLK3 | 77 |
| THAI11 | PLK4 | PLK4 | 100 |
| THAI11 | PRKCD | PRKCD | 94 |
| THAI11 | PRKCE | PRKCE | 74 |
| THAI11 | PRKCH | PRKCH | 100 |
| THAI11 | PRKCI | PRKCI | 100 |
| THAI11 | PRKCQ | PRKCQ | 90 |
| THAI11 | PRKD1 | PRKD1 | 100 |
| THAI11 | PRKD2 | PRKD2 | 86 |
| THAI11 | PRKD3 | PRKD3 | 68 |
| THAI11 | PRKG1 | PRKG1 | 100 |
| THAI11 | PRKG2 | PRKG2 | 86 |
| THAI11 | PRKR | EIF2AK2 | 100 |
| THAI11 | PRKX | PRKX | 100 |
| THAI11 | PRP4 | PRPF4B | 96 |
| THAI11 | PYK2 | PTK2B | 85 |
| THAI11 | QSK | KIAA0999 | 75 |
| THAI11 | RAF1 | RAF1 | 96 |
| THAI11 | RET | RET | 100 |
| THAI11 | RET(M918T) | RET | 79 |
| THAI11 | RET(V804L) | RET | 81 |
| THAI11 | RET(V804M) | RET | 100 |
| THAI11 | RIOK1 | RIOK1 | 74 |
| THAI11 | RIOK2 | RIOK2 | 86 |
| THAI11 | RIOK3 | RIOK3 | 75 |
| THAI11 | RIPK1 | RIPK1 | 98 |
| THAI11 | RIPK2 | RIPK2 | 68 |
| THAI11 | RIPK4 | RIPK4 | 92 |
| THAI11 | RIPK5 | DSTYK | 68 |
| THAI11 | ROCK1 | ROCK1 | 100 |
| THAI11 | ROCK2 | ROCK2 | 100 |
| THAI11 | ROS1 | ROS1 | 76 |
| THAI11 | RPS6KA4(Kin.Dom.1-N-terminal) | RPS6KA4 | 95 |
| THAI11 | RPS6KA4(Kin.Dom.2-C-terminal) | RPS6KA4 | 86 |
| THAI11 | RPS6KA5(Kin.Dom.1-N-terminal) | RPS6KA5 | 100 |

| | | | |
|---|---|---|---|
| THAI11 | RPS6KA5(Kin.Dom.2-C-terminal) | RPS6KA5 | 74 |
| THAI11 | RSK1(Kin.Dom.1-N-terminal) | RPS6KA1 | 100 |
| THAI11 | RSK1(Kin.Dom.2-C-terminal) | RPS6KA1 | 99 |
| THAI11 | RSK2(Kin.Dom.1-N-terminal) | RPS6KA3 | 93 |
| THAI11 | RSK2(Kin.Dom.2-C-terminal) | RPS6KA3 | 100 |
| THAI11 | RSK3(Kin.Dom.1-N-terminal) | RPS6KA2 | 100 |
| THAI11 | RSK3(Kin.Dom.2-C-terminal) | RPS6KA2 | 100 |
| THAI11 | RSK4(Kin.Dom.1-N-terminal) | RPS6KA6 | 89 |
| THAI11 | RSK4(Kin.Dom.2-C-terminal) | RPS6KA6 | 95 |
| THAI11 | S6K1 | RPS6KB1 | 96 |
| THAI11 | SBK1 | SBK1 | 56 |
| THAI11 | SGK | SGK1 | 85 |
| THAI11 | SgK110 | SgK110 | 56 |
| THAI11 | SGK2 | SGK2 | 74 |
| THAI11 | SGK3 | SGK3 | 93 |
| THAI11 | SIK | SIK1 | 84 |
| THAI11 | SIK2 | SIK2 | 100 |
| THAI11 | SLK | SLK | 70 |
| THAI11 | SNARK | NUAK2 | 85 |
| THAI11 | SNRK | SNRK | 100 |
| THAI11 | SRC | SRC | 91 |
| THAI11 | SRMS | SRMS | 81 |
| THAI11 | SRPK1 | SRPK1 | 100 |
| THAI11 | SRPK2 | SRPK2 | 98 |
| THAI11 | SRPK3 | SRPK3 | 84 |
| THAI11 | STK16 | STK16 | 71 |
| THAI11 | STK33 | STK33 | 100 |
| THAI11 | STK35 | STK35 | 97 |
| THAI11 | STK36 | STK36 | 86 |
| THAI11 | STK39 | STK39 | 75 |
| THAI11 | SYK | SYK | 81 |
| THAI11 | TAK1 | MAP3K7 | 83 |
| THAI11 | TAOK1 | TAOK1 | 100 |
| THAI11 | TAOK2 | TAOK2 | 100 |
| THAI11 | TAOK3 | TAOK3 | 54 |
| THAI11 | TBK1 | TBK1 | 73 |
| THAI11 | TEC | TEC | 100 |
| THAI11 | TESK1 | TESK1 | 94 |
| THAI11 | TGFBR1 | TGFBR1 | 85 |
| THAI11 | TGFBR2 | TGFBR2 | 89 |
| THAI11 | TIE1 | TIE1 | 100 |
| THAI11 | TIE2 | TEK | 99 |
| THAI11 | TLK1 | TLK1 | 84 |
| THAI11 | TLK2 | TLK2 | 100 |

| | | | |
|---|---|---|---|
| THAI11 | TNIK | TNIK | 84 |
| THAI11 | TNK1 | TNK1 | 81 |
| THAI11 | TNK2 | TNK2 | 79 |
| THAI11 | TNNI3K | TNNI3K | 51 |
| THAI11 | TRKA | NTRK1 | 100 |
| THAI11 | TRKB | NTRK2 | 100 |
| THAI11 | TRKC | NTRK3 | 100 |
| THAI11 | TRPM6 | TRPM6 | 91 |
| THAI11 | TSSK1B | TSSK1B | 98 |
| THAI11 | TSSK3 | TSSK3 | 100 |
| THAI11 | TTK | TTK | 61 |
| THAI11 | TXK | TXK | 88 |
| THAI11 | TYK2(JH1domain-catalytic) | TYK2 | 100 |
| THAI11 | TYK2(JH2domain-pseudokinase) | TYK2 | 83 |
| THAI11 | TYRO3 | TYRO3 | 96 |
| THAI11 | ULK1 | ULK1 | 59 |
| THAI11 | ULK2 | ULK2 | 89 |
| THAI11 | ULK3 | ULK3 | 71 |
| THAI11 | VEGFR2 | KDR | 86 |
| THAI11 | VPS34 | PIK3C3 | 98 |
| THAI11 | VRK2 | VRK2 | 78 |
| THAI11 | WEE1 | WEE1 | 92 |
| THAI11 | WEE2 | WEE2 | 82 |
| THAI11 | WNK1 | WNK1 | 92 |
| THAI11 | WNK2 | WNK2 | 84 |
| THAI11 | WNK3 | WNK3 | 73 |
| THAI11 | WNK4 | WNK4 | 64 |
| THAI11 | YANK1 | STK32A | 100 |
| THAI11 | YANK2 | STK32B | 100 |
| THAI11 | YANK3 | STK32C | 84 |
| THAI11 | YES | YES1 | 72 |
| THAI11 | YSK1 | STK25 | 92 |
| THAI11 | YSK4 | MAP3K19 | 70 |
| THAI11 | ZAK | ZAK | 86 |
| THAI11 | ZAP70 | ZAP70 | 100 |

**Table S7**. Kinome scan of THAI14 (**16**) @ 1 µM against 469 kinases from DiscoverX/Eurofins.

| Compound Name | DiscoveRx Gene Symbol | Entrez Gene Symbol | Percent Control |
|---|---|---|---|
| THAI14 | AAK1 | AAK1 | 79 |
| THAI14 | ABL1(E255K)-phosphorylated | ABL1 | 74 |
| THAI14 | ABL1(F317I)-nonphosphorylated | ABL1 | 68 |
| THAI14 | ABL1(F317I)-phosphorylated | ABL1 | 100 |

| | | | |
|---|---|---|---|
| THAI14 | ABL1(F317L)-nonphosphorylated | ABL1 | 50 |
| THAI14 | ABL1(F317L)-phosphorylated | ABL1 | 87 |
| THAI14 | ABL1(H396P)-nonphosphorylated | ABL1 | 69 |
| THAI14 | ABL1(H396P)-phosphorylated | ABL1 | 64 |
| THAI14 | ABL1(M351T)-phosphorylated | ABL1 | 86 |
| THAI14 | ABL1(Q252H)-nonphosphorylated | ABL1 | 70 |
| THAI14 | ABL1(Q252H)-phosphorylated | ABL1 | 74 |
| THAI14 | ABL1(T315I)-nonphosphorylated | ABL1 | 73 |
| THAI14 | ABL1(T315I)-phosphorylated | ABL1 | 89 |
| THAI14 | ABL1(Y253F)-phosphorylated | ABL1 | 71 |
| THAI14 | ABL1-nonphosphorylated | ABL1 | 55 |
| THAI14 | ABL1-phosphorylated | ABL1 | 56 |
| THAI14 | ABL2 | ABL2 | 91 |
| THAI14 | ACVR1 | ACVR1 | 57 |
| THAI14 | ACVR1B | ACVR1B | 82 |
| THAI14 | ACVR2A | ACVR2A | 94 |
| THAI14 | ACVR2B | ACVR2B | 96 |
| THAI14 | ACVRL1 | ACVRL1 | 56 |
| THAI14 | ADCK3 | CABC1 | 100 |
| THAI14 | ADCK4 | ADCK4 | 78 |
| THAI14 | AKT1 | AKT1 | 95 |
| THAI14 | AKT2 | AKT2 | 78 |
| THAI14 | AKT3 | AKT3 | 91 |
| THAI14 | ALK | ALK | 87 |
| THAI14 | ALK(C1156Y) | ALK | 90 |
| THAI14 | ALK(L1196M) | ALK | 78 |
| THAI14 | AMPK-alpha1 | PRKAA1 | 84 |
| THAI14 | AMPK-alpha2 | PRKAA2 | 100 |
| THAI14 | ANKK1 | ANKK1 | 95 |
| THAI14 | ARK5 | NUAK1 | 100 |
| THAI14 | ASK1 | MAP3K5 | 87 |
| THAI14 | ASK2 | MAP3K6 | 100 |
| THAI14 | AURKA | AURKA | 88 |
| THAI14 | AURKB | AURKB | 85 |
| THAI14 | AURKC | AURKC | 68 |
| THAI14 | AXL | AXL | 79 |
| THAI14 | BIKE | BMP2K | 68 |
| THAI14 | BLK | BLK | 93 |
| THAI14 | BMPR1A | BMPR1A | 67 |
| THAI14 | BMPR1B | BMPR1B | 79 |
| THAI14 | BMPR2 | BMPR2 | 86 |
| THAI14 | BMX | BMX | 80 |
| THAI14 | BRAF | BRAF | 60 |
| THAI14 | BRAF(V600E) | BRAF | 75 |

| | | | |
|---|---|---|---|
| THAI14 | BRK | PTK6 | 68 |
| THAI14 | BRSK1 | BRSK1 | 97 |
| THAI14 | BRSK2 | BRSK2 | 100 |
| THAI14 | BTK | BTK | 60 |
| THAI14 | BUB1 | BUB1 | 72 |
| THAI14 | CAMK1 | CAMK1 | 100 |
| THAI14 | CAMK1B | PNCK | 71 |
| THAI14 | CAMK1D | CAMK1D | 100 |
| THAI14 | CAMK1G | CAMK1G | 88 |
| THAI14 | CAMK2A | CAMK2A | 77 |
| THAI14 | CAMK2B | CAMK2B | 99 |
| THAI14 | CAMK2D | CAMK2D | 74 |
| THAI14 | CAMK2G | CAMK2G | 86 |
| THAI14 | CAMK4 | CAMK4 | 50 |
| THAI14 | CAMKK1 | CAMKK1 | 85 |
| THAI14 | CAMKK2 | CAMKK2 | 76 |
| THAI14 | CASK | CASK | 82 |
| THAI14 | CDC2L1 | CDK11B | 87 |
| THAI14 | CDC2L2 | CDC2L2 | 100 |
| THAI14 | CDC2L5 | CDK13 | 97 |
| THAI14 | CDK11 | CDK19 | 100 |
| THAI14 | CDK2 | CDK2 | 100 |
| THAI14 | CDK3 | CDK3 | 89 |
| THAI14 | CDK4 | CDK4 | 100 |
| THAI14 | CDK4-cyclinD1 | CDK4 | 73 |
| THAI14 | CDK4-cyclinD3 | CDK4 | 69 |
| THAI14 | CDK5 | CDK5 | 100 |
| THAI14 | CDK7 | CDK7 | 74 |
| THAI14 | CDK8 | CDK8 | 100 |
| THAI14 | CDK9 | CDK9 | 89 |
| THAI14 | CDKL1 | CDKL1 | 68 |
| THAI14 | CDKL2 | CDKL2 | 99 |
| THAI14 | CDKL3 | CDKL3 | 87 |
| THAI14 | CDKL5 | CDKL5 | 86 |
| THAI14 | CHEK1 | CHEK1 | 80 |
| THAI14 | CHEK2 | CHEK2 | 100 |
| THAI14 | CIT | CIT | 81 |
| THAI14 | CLK1 | CLK1 | 86 |
| THAI14 | CLK2 | CLK2 | 79 |
| THAI14 | CLK3 | CLK3 | 73 |
| THAI14 | CLK4 | CLK4 | 100 |
| THAI14 | CSF1R | CSF1R | 79 |
| THAI14 | CSF1R-autoinhibited | CSF1R | 81 |
| THAI14 | CSK | CSK | 100 |

| | | | |
|---|---|---|---|
| THAI14 | CSNK1A1 | CSNK1A1 | 70 |
| THAI14 | CSNK1A1L | CSNK1A1L | 85 |
| THAI14 | CSNK1D | CSNK1D | 99 |
| THAI14 | CSNK1E | CSNK1E | 97 |
| THAI14 | CSNK1G1 | CSNK1G1 | 87 |
| THAI14 | CSNK1G2 | CSNK1G2 | 90 |
| THAI14 | CSNK1G3 | CSNK1G3 | 100 |
| THAI14 | CSNK2A1 | CSNK2A1 | 65 |
| THAI14 | CSNK2A2 | CSNK2A2 | 94 |
| THAI14 | CTK | MATK | 86 |
| THAI14 | DAPK1 | DAPK1 | 91 |
| THAI14 | DAPK2 | DAPK2 | 93 |
| THAI14 | DAPK3 | DAPK3 | 80 |
| THAI14 | DCAMKL1 | DCLK1 | 63 |
| THAI14 | DCAMKL2 | DCLK2 | 95 |
| THAI14 | DCAMKL3 | DCLK3 | 69 |
| THAI14 | DDR1 | DDR1 | 80 |
| THAI14 | DDR2 | DDR2 | 88 |
| THAI14 | DLK | MAP3K12 | 81 |
| THAI14 | DMPK | DMPK | 76 |
| THAI14 | DMPK2 | CDC42BPG | 81 |
| THAI14 | DRAK1 | STK17A | 100 |
| THAI14 | DRAK2 | STK17B | 98 |
| THAI14 | DYRK1A | DYRK1A | 81 |
| THAI14 | DYRK1B | DYRK1B | 100 |
| THAI14 | DYRK2 | DYRK2 | 82 |
| THAI14 | EGFR | EGFR | 93 |
| THAI14 | EGFR(E746-A750del) | EGFR | 89 |
| THAI14 | EGFR(G719C) | EGFR | 80 |
| THAI14 | EGFR(G719S) | EGFR | 100 |
| THAI14 | EGFR(L747-E749del, A750P) | EGFR | 99 |
| THAI14 | EGFR(L747-S752del, P753S) | EGFR | 82 |
| THAI14 | EGFR(L747-T751del,Sins) | EGFR | 100 |
| THAI14 | EGFR(L858R) | EGFR | 82 |
| THAI14 | EGFR(L858R,T790M) | EGFR | 77 |
| THAI14 | EGFR(L861Q) | EGFR | 81 |
| THAI14 | EGFR(S752-I759del) | EGFR | 88 |
| THAI14 | EGFR(T790M) | EGFR | 72 |
| THAI14 | EIF2AK1 | EIF2AK1 | 64 |
| THAI14 | EPHA1 | EPHA1 | 82 |
| THAI14 | EPHA2 | EPHA2 | 81 |
| THAI14 | EPHA3 | EPHA3 | 100 |
| THAI14 | EPHA4 | EPHA4 | 94 |
| THAI14 | EPHA5 | EPHA5 | 90 |

| | | | |
|---|---|---|---|
| THAI14 | EPHA6 | EPHA6 | 100 |
| THAI14 | EPHA7 | EPHA7 | 79 |
| THAI14 | EPHA8 | EPHA8 | 82 |
| THAI14 | EPHB1 | EPHB1 | 96 |
| THAI14 | EPHB2 | EPHB2 | 80 |
| THAI14 | EPHB3 | EPHB3 | 100 |
| THAI14 | EPHB4 | EPHB4 | 91 |
| THAI14 | EPHB6 | EPHB6 | 75 |
| THAI14 | ERBB2 | ERBB2 | 89 |
| THAI14 | ERBB3 | ERBB3 | 75 |
| THAI14 | ERBB4 | ERBB4 | 91 |
| THAI14 | ERK1 | MAPK3 | 86 |
| THAI14 | ERK2 | MAPK1 | 91 |
| THAI14 | ERK3 | MAPK6 | 80 |
| THAI14 | ERK4 | MAPK4 | 91 |
| THAI14 | ERK5 | MAPK7 | 84 |
| THAI14 | ERK8 | MAPK15 | 100 |
| THAI14 | ERN1 | ERN1 | 77 |
| THAI14 | FAK | PTK2 | 75 |
| THAI14 | FER | FER | 68 |
| THAI14 | FES | FES | 74 |
| THAI14 | FGFR1 | FGFR1 | 100 |
| THAI14 | FGFR2 | FGFR2 | 90 |
| THAI14 | FGFR3 | FGFR3 | 80 |
| THAI14 | FGFR3(G697C) | FGFR3 | 78 |
| THAI14 | FGFR4 | FGFR4 | 72 |
| THAI14 | FGR | FGR | 83 |
| THAI14 | FLT1 | FLT1 | 88 |
| THAI14 | FLT3 | FLT3 | 89 |
| THAI14 | FLT3(D835H) | FLT3 | 62 |
| THAI14 | FLT3(D835V) | FLT3 | 84 |
| THAI14 | FLT3(D835Y) | FLT3 | 72 |
| THAI14 | FLT3(ITD) | FLT3 | 82 |
| THAI14 | FLT3(ITD,D835V) | FLT3 | 64 |
| THAI14 | FLT3(ITD,F691L) | FLT3 | 70 |
| THAI14 | FLT3(K663Q) | FLT3 | 85 |
| THAI14 | FLT3(N841I) | FLT3 | 82 |
| THAI14 | FLT3(R834Q) | FLT3 | 100 |
| THAI14 | FLT3-autoinhibited | FLT3 | 90 |
| THAI14 | FLT4 | FLT4 | 64 |
| THAI14 | FRK | FRK | 70 |
| THAI14 | FYN | FYN | 64 |
| THAI14 | GAK | GAK | 89 |
| THAI14 | GCN2(Kin.Dom.2,S808G) | EIF2AK4 | 96 |

| | | | |
|---|---|---|---|
| THAI14 | GRK1 | GRK1 | 59 |
| THAI14 | GRK2 | ADRBK1 | 71 |
| THAI14 | GRK3 | ADRBK2 | 66 |
| THAI14 | GRK4 | GRK4 | 93 |
| THAI14 | GRK7 | GRK7 | 78 |
| THAI14 | GSK3A | GSK3A | 88 |
| THAI14 | GSK3B | GSK3B | 69 |
| THAI14 | HASPIN | GSG2 | 76 |
| THAI14 | HCK | HCK | 95 |
| THAI14 | HIPK1 | HIPK1 | 55 |
| THAI14 | HIPK2 | HIPK2 | 100 |
| THAI14 | HIPK3 | HIPK3 | 85 |
| THAI14 | HIPK4 | HIPK4 | 100 |
| THAI14 | HPK1 | MAP4K1 | 97 |
| THAI14 | HUNK | HUNK | 73 |
| THAI14 | ICK | ICK | 86 |
| THAI14 | IGF1R | IGF1R | 99 |
| THAI14 | IKK-alpha | CHUK | 72 |
| THAI14 | IKK-beta | IKBKB | 82 |
| THAI14 | IKK-epsilon | IKBKE | 90 |
| THAI14 | INSR | INSR | 66 |
| THAI14 | INSRR | INSRR | 98 |
| THAI14 | IRAK1 | IRAK1 | 90 |
| THAI14 | IRAK3 | IRAK3 | 100 |
| THAI14 | IRAK4 | IRAK4 | 62 |
| THAI14 | ITK | ITK | 86 |
| THAI14 | JAK1(JH1domain-catalytic) | JAK1 | 94 |
| THAI14 | JAK1(JH2domain-pseudokinase) | JAK1 | 100 |
| THAI14 | JAK2(JH1domain-catalytic) | JAK2 | 74 |
| THAI14 | JAK3(JH1domain-catalytic) | JAK3 | 72 |
| THAI14 | JNK1 | MAPK8 | 72 |
| THAI14 | JNK2 | MAPK9 | 67 |
| THAI14 | JNK3 | MAPK10 | 76 |
| THAI14 | KIT | KIT | 100 |
| THAI14 | KIT(A829P) | KIT | 70 |
| THAI14 | KIT(D816H) | KIT | 80 |
| THAI14 | KIT(D816V) | KIT | 76 |
| THAI14 | KIT(L576P) | KIT | 89 |
| THAI14 | KIT(V559D) | KIT | 99 |
| THAI14 | KIT(V559D,T670I) | KIT | 77 |
| THAI14 | KIT(V559D,V654A) | KIT | 61 |
| THAI14 | KIT-autoinhibited | KIT | 77 |
| THAI14 | LATS1 | LATS1 | 97 |
| THAI14 | LATS2 | LATS2 | 76 |

| | | | |
|---|---|---|---|
| THAI14 | LCK | LCK | 98 |
| THAI14 | LIMK1 | LIMK1 | 68 |
| THAI14 | LIMK2 | LIMK2 | 76 |
| THAI14 | LKB1 | STK11 | 100 |
| THAI14 | LOK | STK10 | 81 |
| THAI14 | LRRK2 | LRRK2 | 100 |
| THAI14 | LRRK2(G2019S) | LRRK2 | 100 |
| THAI14 | LTK | LTK | 75 |
| THAI14 | LYN | LYN | 100 |
| THAI14 | LZK | MAP3K13 | 93 |
| THAI14 | MAK | MAK | 87 |
| THAI14 | MAP3K1 | MAP3K1 | 73 |
| THAI14 | MAP3K15 | MAP3K15 | 82 |
| THAI14 | MAP3K2 | MAP3K2 | 89 |
| THAI14 | MAP3K3 | MAP3K3 | 62 |
| THAI14 | MAP3K4 | MAP3K4 | 77 |
| THAI14 | MAP4K2 | MAP4K2 | 100 |
| THAI14 | MAP4K3 | MAP4K3 | 92 |
| THAI14 | MAP4K4 | MAP4K4 | 100 |
| THAI14 | MAP4K5 | MAP4K5 | 100 |
| THAI14 | MAPKAPK2 | MAPKAPK2 | 92 |
| THAI14 | MAPKAPK5 | MAPKAPK5 | 100 |
| THAI14 | MARK1 | MARK1 | 100 |
| THAI14 | MARK2 | MARK2 | 100 |
| THAI14 | MARK3 | MARK3 | 100 |
| THAI14 | MARK4 | MARK4 | 94 |
| THAI14 | MAST1 | MAST1 | 100 |
| THAI14 | MEK1 | MAP2K1 | 92 |
| THAI14 | MEK2 | MAP2K2 | 82 |
| THAI14 | MEK3 | MAP2K3 | 83 |
| THAI14 | MEK4 | MAP2K4 | 86 |
| THAI14 | MEK5 | MAP2K5 | 80 |
| THAI14 | MEK6 | MAP2K6 | 100 |
| THAI14 | MELK | MELK | 100 |
| THAI14 | MERTK | MERTK | 100 |
| THAI14 | MET | MET | 83 |
| THAI14 | MET(M1250T) | MET | 99 |
| THAI14 | MET(Y1235D) | MET | 87 |
| THAI14 | MINK | MINK1 | 83 |
| THAI14 | MKK7 | MAP2K7 | 89 |
| THAI14 | MKNK1 | MKNK1 | 93 |
| THAI14 | MKNK2 | MKNK2 | 60 |
| THAI14 | MLCK | MYLK3 | 83 |
| THAI14 | MLK1 | MAP3K9 | 83 |

| | | | |
|---|---|---|---|
| THAI14 | MLK2 | MAP3K10 | 100 |
| THAI14 | MLK3 | MAP3K11 | 88 |
| THAI14 | MRCKA | CDC42BPA | 92 |
| THAI14 | MRCKB | CDC42BPB | 79 |
| THAI14 | MST1 | STK4 | 73 |
| THAI14 | MST1R | MST1R | 100 |
| THAI14 | MST2 | STK3 | 92 |
| THAI14 | MST3 | STK24 | 77 |
| THAI14 | MST4 | MST4 | 96 |
| THAI14 | MTOR | MTOR | 81 |
| THAI14 | MUSK | MUSK | 86 |
| THAI14 | MYLK | MYLK | 61 |
| THAI14 | MYLK2 | MYLK2 | 77 |
| THAI14 | MYLK4 | MYLK4 | 88 |
| THAI14 | MYO3A | MYO3A | 83 |
| THAI14 | MYO3B | MYO3B | 94 |
| THAI14 | NDR1 | STK38 | 79 |
| THAI14 | NDR2 | STK38L | 79 |
| THAI14 | NEK1 | NEK1 | 100 |
| THAI14 | NEK10 | NEK10 | 79 |
| THAI14 | NEK11 | NEK11 | 100 |
| THAI14 | NEK2 | NEK2 | 100 |
| THAI14 | NEK3 | NEK3 | 99 |
| THAI14 | NEK4 | NEK4 | 88 |
| THAI14 | NEK5 | NEK5 | 0 |
| THAI14 | NEK6 | NEK6 | 88 |
| THAI14 | NEK7 | NEK7 | 73 |
| THAI14 | NEK9 | NEK9 | 71 |
| THAI14 | NIK | MAP3K14 | 89 |
| THAI14 | NIM1 | MGC42105 | 90 |
| THAI14 | NLK | NLK | 86 |
| THAI14 | OSR1 | OXSR1 | 100 |
| THAI14 | p38-alpha | MAPK14 | 100 |
| THAI14 | p38-beta | MAPK11 | 75 |
| THAI14 | p38-delta | MAPK13 | 94 |
| THAI14 | p38-gamma | MAPK12 | 79 |
| THAI14 | PAK1 | PAK1 | 86 |
| THAI14 | PAK2 | PAK2 | 100 |
| THAI14 | PAK3 | PAK3 | 77 |
| THAI14 | PAK4 | PAK4 | 100 |
| THAI14 | PAK6 | PAK6 | 100 |
| THAI14 | PAK7 | PAK7 | 100 |
| THAI14 | PCTK1 | CDK16 | 92 |
| THAI14 | PCTK2 | CDK17 | 88 |

| | | | |
|---|---|---|---|
| THAI14 | PCTK3 | CDK18 | 82 |
| THAI14 | PDGFRA | PDGFRA | 100 |
| THAI14 | PDGFRB | PDGFRB | 93 |
| THAI14 | PDPK1 | PDPK1 | 100 |
| THAI14 | PFCDPK1(P.falciparum) | CDPK1 | 79 |
| THAI14 | PFPK5(P.falciparum) | MAL13P1.279 | 89 |
| THAI14 | PFTAIRE2 | CDK15 | 70 |
| THAI14 | PFTK1 | CDK14 | 95 |
| THAI14 | PHKG1 | PHKG1 | 92 |
| THAI14 | PHKG2 | PHKG2 | 85 |
| THAI14 | PIK3C2B | PIK3C2B | 86 |
| THAI14 | PIK3C2G | PIK3C2G | 79 |
| THAI14 | PIK3CA | PIK3CA | 100 |
| THAI14 | PIK3CA(C420R) | PIK3CA | 63 |
| THAI14 | PIK3CA(E542K) | PIK3CA | 38 |
| THAI14 | PIK3CA(E545A) | PIK3CA | 87 |
| THAI14 | PIK3CA(E545K) | PIK3CA | 89 |
| THAI14 | PIK3CA(H1047L) | PIK3CA | 100 |
| THAI14 | PIK3CA(H1047Y) | PIK3CA | 100 |
| THAI14 | PIK3CA(I800L) | PIK3CA | 89 |
| THAI14 | PIK3CA(M1043I) | PIK3CA | 100 |
| THAI14 | PIK3CA(Q546K) | PIK3CA | 46 |
| THAI14 | PIK3CB | PIK3CB | 100 |
| THAI14 | PIK3CD | PIK3CD | 85 |
| THAI14 | PIK3CG | PIK3CG | 100 |
| THAI14 | PIK4CB | PI4KB | 91 |
| THAI14 | PIKFYVE | PIKFYVE | 81 |
| THAI14 | PIM1 | PIM1 | 100 |
| THAI14 | PIM2 | PIM2 | 97 |
| THAI14 | PIM3 | PIM3 | 100 |
| THAI14 | PIP5K1A | PIP5K1A | 100 |
| THAI14 | PIP5K1C | PIP5K1C | 80 |
| THAI14 | PIP5K2B | PIP4K2B | 96 |
| THAI14 | PIP5K2C | PIP4K2C | 99 |
| THAI14 | PKAC-alpha | PRKACA | 98 |
| THAI14 | PKAC-beta | PRKACB | 88 |
| THAI14 | PKMYT1 | PKMYT1 | 94 |
| THAI14 | PKN1 | PKN1 | 91 |
| THAI14 | PKN2 | PKN2 | 95 |
| THAI14 | PKNB(M.tuberculosis) | pknB | 89 |
| THAI14 | PLK1 | PLK1 | 97 |
| THAI14 | PLK2 | PLK2 | 87 |
| THAI14 | PLK3 | PLK3 | 78 |
| THAI14 | PLK4 | PLK4 | 95 |

| | | | |
|---|---|---|---|
| THAI14 | PRKCD | PRKCD | 100 |
| THAI14 | PRKCE | PRKCE | 71 |
| THAI14 | PRKCH | PRKCH | 100 |
| THAI14 | PRKCI | PRKCI | 86 |
| THAI14 | PRKCQ | PRKCQ | 71 |
| THAI14 | PRKD1 | PRKD1 | 62 |
| THAI14 | PRKD2 | PRKD2 | 83 |
| THAI14 | PRKD3 | PRKD3 | 62 |
| THAI14 | PRKG1 | PRKG1 | 86 |
| THAI14 | PRKG2 | PRKG2 | 66 |
| THAI14 | PRKR | EIF2AK2 | 100 |
| THAI14 | PRKX | PRKX | 96 |
| THAI14 | PRP4 | PRPF4B | 100 |
| THAI14 | PYK2 | PTK2B | 83 |
| THAI14 | QSK | KIAA0999 | 80 |
| THAI14 | RAF1 | RAF1 | 88 |
| THAI14 | RET | RET | 93 |
| THAI14 | RET(M918T) | RET | 77 |
| THAI14 | RET(V804L) | RET | 88 |
| THAI14 | RET(V804M) | RET | 77 |
| THAI14 | RIOK1 | RIOK1 | 69 |
| THAI14 | RIOK2 | RIOK2 | 79 |
| THAI14 | RIOK3 | RIOK3 | 69 |
| THAI14 | RIPK1 | RIPK1 | 84 |
| THAI14 | RIPK2 | RIPK2 | 55 |
| THAI14 | RIPK4 | RIPK4 | 79 |
| THAI14 | RIPK5 | DSTYK | 80 |
| THAI14 | ROCK1 | ROCK1 | 100 |
| THAI14 | ROCK2 | ROCK2 | 100 |
| THAI14 | ROS1 | ROS1 | 75 |
| THAI14 | RPS6KA4(Kin.Dom.1-N-terminal) | RPS6KA4 | 74 |
| THAI14 | RPS6KA4(Kin.Dom.2-C-terminal) | RPS6KA4 | 91 |
| THAI14 | RPS6KA5(Kin.Dom.1-N-terminal) | RPS6KA5 | 100 |
| THAI14 | RPS6KA5(Kin.Dom.2-C-terminal) | RPS6KA5 | 87 |
| THAI14 | RSK1(Kin.Dom.1-N-terminal) | RPS6KA1 | 78 |
| THAI14 | RSK1(Kin.Dom.2-C-terminal) | RPS6KA1 | 86 |
| THAI14 | RSK2(Kin.Dom.1-N-terminal) | RPS6KA3 | 90 |
| THAI14 | RSK2(Kin.Dom.2-C-terminal) | RPS6KA3 | 100 |
| THAI14 | RSK3(Kin.Dom.1-N-terminal) | RPS6KA2 | 100 |
| THAI14 | RSK3(Kin.Dom.2-C-terminal) | RPS6KA2 | 79 |
| THAI14 | RSK4(Kin.Dom.1-N-terminal) | RPS6KA6 | 87 |
| THAI14 | RSK4(Kin.Dom.2-C-terminal) | RPS6KA6 | 76 |
| THAI14 | S6K1 | RPS6KB1 | 89 |
| THAI14 | SBK1 | SBK1 | 100 |

| | | | |
|---|---|---|---|
| THAI14 | SGK | SGK1 | 90 |
| THAI14 | SgK110 | SgK110 | 67 |
| THAI14 | SGK2 | SGK2 | 90 |
| THAI14 | SGK3 | SGK3 | 100 |
| THAI14 | SIK | SIK1 | 97 |
| THAI14 | SIK2 | SIK2 | 76 |
| THAI14 | SLK | SLK | 77 |
| THAI14 | SNARK | NUAK2 | 93 |
| THAI14 | SNRK | SNRK | 77 |
| THAI14 | SRC | SRC | 100 |
| THAI14 | SRMS | SRMS | 83 |
| THAI14 | SRPK1 | SRPK1 | 68 |
| THAI14 | SRPK2 | SRPK2 | 94 |
| THAI14 | SRPK3 | SRPK3 | 92 |
| THAI14 | STK16 | STK16 | 89 |
| THAI14 | STK33 | STK33 | 100 |
| THAI14 | STK35 | STK35 | 83 |
| THAI14 | STK36 | STK36 | 100 |
| THAI14 | STK39 | STK39 | 70 |
| THAI14 | SYK | SYK | 98 |
| THAI14 | TAK1 | MAP3K7 | 80 |
| THAI14 | TAOK1 | TAOK1 | 88 |
| THAI14 | TAOK2 | TAOK2 | 77 |
| THAI14 | TAOK3 | TAOK3 | 39 |
| THAI14 | TBK1 | TBK1 | 58 |
| THAI14 | TEC | TEC | 82 |
| THAI14 | TESK1 | TESK1 | 89 |
| THAI14 | TGFBR1 | TGFBR1 | 52 |
| THAI14 | TGFBR2 | TGFBR2 | 59 |
| THAI14 | TIE1 | TIE1 | 92 |
| THAI14 | TIE2 | TEK | 81 |
| THAI14 | TLK1 | TLK1 | 79 |
| THAI14 | TLK2 | TLK2 | 100 |
| THAI14 | TNIK | TNIK | 75 |
| THAI14 | TNK1 | TNK1 | 55 |
| THAI14 | TNK2 | TNK2 | 58 |
| THAI14 | TNNI3K | TNNI3K | 56 |
| THAI14 | TRKA | NTRK1 | 100 |
| THAI14 | TRKB | NTRK2 | 100 |
| THAI14 | TRKC | NTRK3 | 100 |
| THAI14 | TRPM6 | TRPM6 | 100 |
| THAI14 | TSSK1B | TSSK1B | 93 |
| THAI14 | TSSK3 | TSSK3 | 93 |
| THAI14 | TTK | TTK | 87 |

| | | | |
|---|---|---|---|
| THAI14 | TXK | TXK | 79 |
| THAI14 | TYK2(JH1domain-catalytic) | TYK2 | 100 |
| THAI14 | TYK2(JH2domain-pseudokinase) | TYK2 | 96 |
| THAI14 | TYRO3 | TYRO3 | 73 |
| THAI14 | ULK1 | ULK1 | 74 |
| THAI14 | ULK2 | ULK2 | 82 |
| THAI14 | ULK3 | ULK3 | 68 |
| THAI14 | VEGFR2 | KDR | 92 |
| THAI14 | VPS34 | PIK3C3 | 98 |
| THAI14 | VRK2 | VRK2 | 100 |
| THAI14 | WEE1 | WEE1 | 100 |
| THAI14 | WEE2 | WEE2 | 71 |
| THAI14 | WNK1 | WNK1 | 79 |
| THAI14 | WNK2 | WNK2 | 84 |
| THAI14 | WNK3 | WNK3 | 85 |
| THAI14 | WNK4 | WNK4 | 63 |
| THAI14 | YANK1 | STK32A | 77 |
| THAI14 | YANK2 | STK32B | 100 |
| THAI14 | YANK3 | STK32C | 78 |
| THAI14 | YES | YES1 | 55 |
| THAI14 | YSK1 | STK25 | 100 |
| THAI14 | YSK4 | MAP3K19 | 83 |
| THAI14 | ZAK | ZAK | 80 |
| THAI14 | ZAP70 | ZAP70 | 100 |

**Table S8**. NanoBRET$^{Tm}$ target engagement assays of compound **5**, **15** and **16** on MAPK9, DDR1, JAK1, PNCK and PIK3CA.

| Target ID | NLuc location | Catalog No. Vector (Promega) | CAS No. Vector (Promega) | Tracer ID (Promega) | Catalog No. Assay (Promega) | CAS No. Assay (Promega) | [Tracer], [nM] |
|---|---|---|---|---|---|---|---|
| MAPK9 | N | NV1711 | N/A | K-5 | N2500 | N/A | 125 |
| DDR1 | C | N2451 | N/A | K-4 | N2520 | N/A | 50 |
| JAK1 | C | N/A | CS1810C386 | K-10 | N2640 | CS1810C122 | 875 |
| PNCK | C | N/A | Kind gift of Promega | K-10 | N2640 | CS1810C122 | 850 |
| PIK3CA (activated with co-transfected PIK3R1) | N | NV3901 (NV4031) | CS1810C52 (CS1810C53) | K-3 | N2600 | CS181016 | 100 |

#### II Radioactive kinase Assay from reaction biology

4 compounds tested against 2 kinases
Compounds were tested in 10-dose IC50 mode with a 3-fold serial dilution starting at 10 μM.
Control compound, LDN193189, was tested in 10-dose IC50 mode with 3-fold serial dilution starting at 10 μM.
Reactions were carried out at 1 μM ATP.
Data pages include raw data, % Enzyme activity (relative to DMSO controls) and curve fits.
*Curve fits were performed where the enzyme activities at the highest concentration of compounds were less than 65%.

#### III Chemistry: Synthetic procedures and characterization of compound 9–16

Synthesis of compound **9**; 2-(5-chloro-2-fluorophenyl)-5-methyl-*N*-(2-nitrophenyl)pyridin-4-amine:

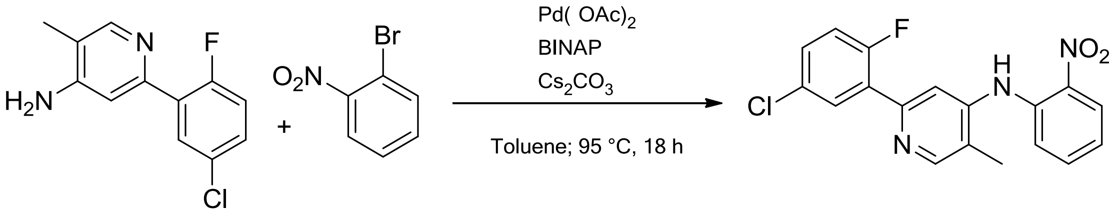

**6** **7** **9**

A mixture of 2-(5-Chloro-2-fluorophenyl)-5-methylpyridin-4-amine (500 mg, 2.11 mmol, 1 equiv), 1-bromo-2-nitrobenzene (852 mg, 4.22 mmol, 2 equiv), $Pd(OAc)_2$ (19 mg, 0.08 mmol, 0.4 equiv), BINAP (53 mg, 0.08 mmol, 0.4 equiv) and $Cs_2CO_3$ (962 mg, 2.95 mmol, 1 equiv) were dissolved in 25 mL toluene and the reaction stirred overnight (18 h) at 95 °C under argon atmosphere. The reaction was cooled to room temperature and the solvent was evaporated under reduced pressure. The crude product was purified by column chromatography on silica gel (*n*-hexane/EtOAc) to obtain 2-(5-chloro-2-fluorophenyl)-5-methyl-*N*-(2-nitrophenyl)pyridin-4-amine as an orange solid. **Yield:** 556 mg (1.54 mmol, 73 %). **$R_f$-Value TLC:** 0.44 (*n*-Hexane/EtOAc 5:1). **$^1$H NMR (500 MHz, $(CD_3)_2SO$):** δ = 8.94 (s, 1H), 8.49 (s, 1H), 8.15 (dd, $^3J_{HH}$ = 8.4, $^4J_{HH}$ = 1.5 Hz, 1H), 7.96 (dd, $^3J_{HH}$ = 6.7, $^4J_{HH}$ = 2.8 Hz, 1H), 7.69–7.64 (m, 1H), 7.54–7.49 (m, 2H), 7.44 (dd, $^3J_{HH}$ = 8.4, $^4J_{HH}$ = 1.0 Hz, 1H), 7.35 (dd, $^3J_{HH}$ = 11.0, $^4J_{HH}$ = 8.8 Hz, 1H), 7.18 (ddd, $^3J_{HH}$ = 7.2, $^4J_{HH}$ = 1.2 Hz, 1H), 2.27 (s, 3H). **$^{13}$C NMR (125 MHz, $(CD_3)_2SO$):** δ = 157.47, 151.51, 149.64, 147.24, 138.17, 137.42, 135.54, 130.21 (d, *J* = 9.1 Hz), 129.79, 129.76, 128.67, 128.52 (d, *J* = 13.0 Hz), 126.28, 123.08, 121.66 (d, *J* = 85.8 Hz), 118.47 (d, *J* = 25.1 Hz), 112.40 (d, *J* = 9.6 Hz), 14.41. **MS (ESI+):** m/z = 358.12 $[M+H]^+$.

Synthesis of compound **10**; 2-(5-chloro-2-fluorophenyl)-5-methyl-*N*-(4-methyl-2-nitrophenyl)pyridin-4-amine:

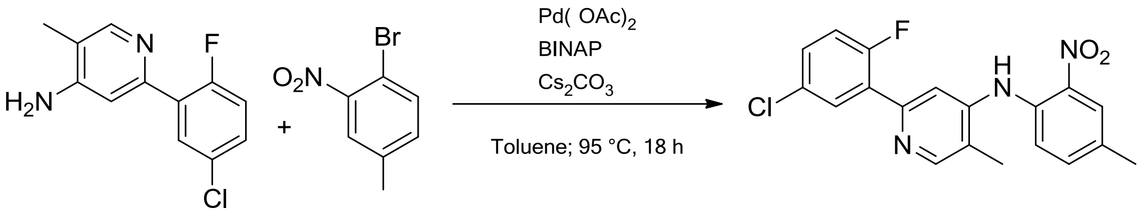

**6** **8** **10**

A mixture of 2-(5-Chloro-2-fluorophenyl)-5-methylpyridin-4-amine (500 mg, 2.11 mmol, 1 equiv), 1-bromo-4-methyl-2-nitrobenzene (912 mg, 4.22 mmol, 2 equiv), $Pd(OAc)_2$ (19 mg, 0.08 mmol, 0.4 equiv), BINAP (53 mg, 0.08 mmol, 0.4 equiv) and $Cs_2CO_3$ (962 mg, 2.95 mmol, 1 equiv) were dissolved in 25 mL toluene and the reaction stirred overnight (18 h) at 95 °C under argon atmosphere. The reaction was cooled to room temperature and the solvent was evaporated under reduced pressure. The crude product was purified by column chromatography on silica gel (*n*-hexane/EtOAc) to obtain 2-(5-chloro-2-fluorophenyl)-5-methyl-*N*-(4-methyl-2-nitrophenyl)pyridin-4-amine as an orange solid. **Yield**: 779 mg (2.10 mmol, 99%). **$R_f$-Value TLC:** 0.48 (*n*-hexane/EtOAc 3:1). **$^1$H NMR (500 MHz, $(CD_3)_2SO$):** δ = 8.69 (s, 1H), 8.43 (s, 1H), 7.99–7.91 (m, 2H), 7.56–7.46 (m, 2H), 7.44–7.37 (m, 2H), 7.34 (dd, $^3J_{HH}$ = 8.8 Hz, 1H), 2.36 (s, 3H), 2.26 (s, 3H). **$^{13}$C NMR (125 MHz, $(CD_3)_2SO$):** δ = 158.44 (d, *J* = 248.4 Hz), 151.24, 149.56, 147.96, 139.26, 136.19, 134.29, 132.73, 130.14 (d, *J* = 9.2 Hz), 129.78 (d, *J* = 3.3 Hz), 128.68 (d, *J* = 3.7 Hz), 128.60, 125.74, 122.81, 121.79, 118.47 (d, *J* = 25.1 Hz), 110.79, 19.90, 14.40. **MS (ESI+):** m/z = 372.14 $[M+H]^+$.

Synthesis of compound **11**; *N*1-[2-(5-chloro-2-fluorophenyl)-5-methylpyridin-4-yl]benzene-1,2-diamine:

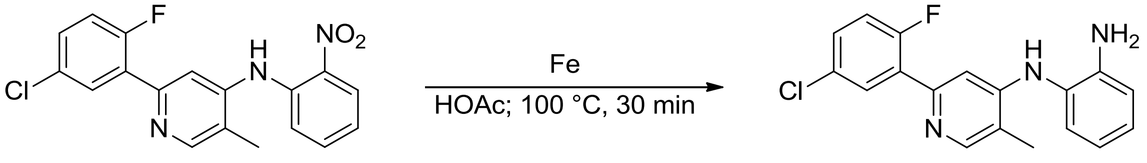

**9** **11**

**9** (556 mg, 1.55 mmol, 1equiv) was dissolved with Fe (608 mg, 10.88 mmol, 7 equiv) in 35 mL HOAc. The reaction stirred at 100 °C for 30 min. The reaction was cooled to RT and 3N NaOH was added to a pH of 10. The product was extracted with EtOAc and the solvent was removed under reduced pressure, to obtain *N*1-[2-(5-chloro-2-fluorophenyl)-5-methylpyridin-4-yl]benzene-1,2-diamine as a green-yellowish solid. **Yield**: 311 mg (0.95 mmol, 61%). **$R_f$-Value TLC:** 0.15 (*n*-hexane/EtOAc 1:1). **$^1$H NMR (250 MHz, $(CD_3)_2SO$):** δ = 8.15 (s, 1H), 7.86 (dd, $^3J_{HH}$ = 6.7, $^4J_{HH}$ = 2.8 Hz), 7.42 (ddd, $^3J_{HH}$ = 4.1, $^4J_{HH}$ = 2.8 Hz), 7.34 (s, 1H), 7.25 (dd, *J* = 8.8 Hz, 1H), 7.03–6.93 (m, 2H), 6.79 (dd, $^3J_{HH}$ = 7.9, $^4J_{HH}$ = 1.4 Hz, 1H), 6.64 (d, $^4J_{HH}$ = 1.6 Hz, 1H), 6.60–6.55 (m, 1H), 4.90 (s, 2H), 2.26 (s, 3H). **MS (ESI+):** m/z = 328.17 $[M+ H]^+$.

Synthesis of compound **12**; *N*1-[2-(5-chloro-2-fluorophenyl)-5-methylpyridin-4-yl]-4-methylbenzene-1,2-diamine:

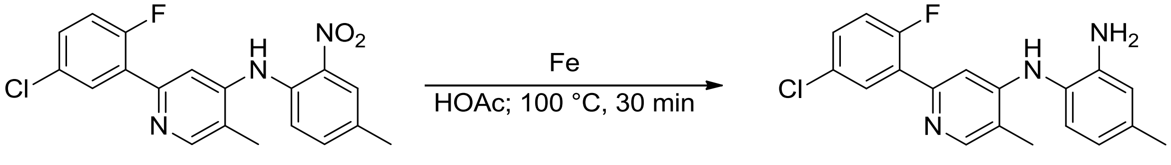

**10** **12**

**10** (727 mg, 1.96 mmol, 1equiv) was dissolved with Fe (764 mg, 13.69 mmol, 7 equiv) in 7 mL HOAc. The reaction stirred at 100 °C for 30 min. The reaction was cooled to RT and 3N NaOH was added to a pH of 10. The product was extracted with EtOAc and the solvent was removed under reduced pressure, to obtain *N*1-[2-(5-chloro-2-fluorophenyl)-5-methylpyridin-4-yl]-4-methylbenzene-1,2-diamine as a yellow solid. **Yield**: 310 mg (0.91 mmol, 47%). **$R_f$-Value TLC:** 0.20 (*n*-hexane/EtOAc 1:1). **$^1$H NMR (500 MHz, $(CD_3)_2SO$):** δ = 8.13 (s, 1H), 7.85 (dd, $^3J_{HH}$ = 6.7, $^4J_{HH}$ = 2.8 Hz, 1H), 7.42 (ddd, $^3J_{HH}$ = 8.7, $^4J_{HH}$ = 2.8 Hz, 1H), 7.28–7.17 (m, 2H), 6.86 (d, $^3J_{HH}$ = 7.8 Hz, 1H), 6.61 (dd, $^3J_{HH}$ = 4.9, $^4J_{HH}$ = 1.8 Hz, 2H), 6.40 (dd, $^3J_{HH}$ = 7.9, $^4J_{HH}$ = 1.9 Hz, 1H), 4.81 (s, 2H), 2.24 (s, 3H), 2.20 (s, 3H). **$^{13}$C NMR (125 MHz, $(CD_3)_2SO$):** δ = 159.18, 157.20, 151.57, 149.55, 148.97 (d, *J* = 2.9 Hz), 144.57, 136.11, 129.70 (d, *J* = 3.7 Hz), 129.50 (d, *J* = 9.6 Hz), 128.40 (d, *J* = 2.9 Hz), 127.81, 121.36, 118.29 (d, *J* = 25.4 Hz), 117.53, 117.22, 115.84, 106.31, 20.97, 14.56 ppm. **MS (ESI+):** m/z = 342.10 $[M+H]^+$.

Synthesis of compound **13**; 1-[2-(5-chloro-2-fluorophenyl)-5-methylpyridin-4-yl]-2,3-dihydro-1*H*-1,3-benzodiazol-2-one:

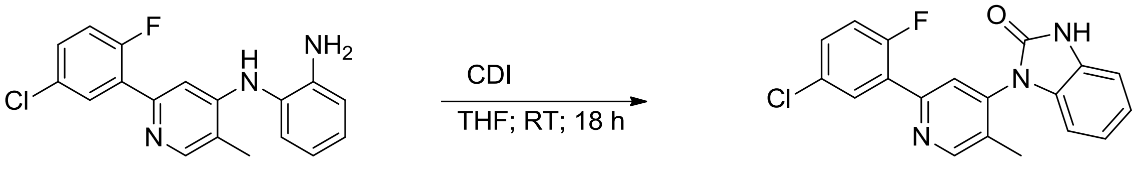

**11** **13**

**11** (311 mg, 0.95 mmol, 1 equiv) was dissolved with CDI (261 mg, 1.61 mmol, 1.7 equiv) in 7 mL THF. The reaction stirred at RT for 18 h under argon atmosphere. The solvent was evaporated under reduced pressure and the crude product was purified by column chromatography on silica gel (*n*-hexane/EtOAc) to obtain 1-[2-(5-chloro-2-fluorophenyl)-5-methylpyridin-4-yl]-2,3-dihydro-1H-1,3-benzodiazol-2-one as a white solid. **Yield:** 245 mg (0.69 mmol, 73%). **$R_f$-Value TLC:** 0.44 (*n*-hexane/EtOAc 1:1). **$^1$H NMR (500 MHz, $(CD_3)_2SO$):** δ = 11.30 (s, 1H), 8.84 (s, 1H), 8.03 (dd, $^3J_{HH}$ = 6.7, $^4J_{HH}$ = 2.8 Hz, 1H), 7.85 (d, $^4J_{HH}$ = 1.5 Hz, 1H), 7.57 (ddd, $^3J_{HH}$ = 4.2, $^4J_{HH}$ = 2.8 Hz, 1H), 7.42 (dd, $^3J_{HH}$ = 8.8 Hz, 1H), 7.11 (d, $^4J_{HH}$ = 1.3 Hz, 2H), 7.01 (ddd, $^3J_{HH}$ = 6.7, $^4J_{HH}$ = 2.0 Hz, 1H), 6.84 (dd, *J* = 7.7, 1.0 Hz, 1H), 2.22 (s, 3H). **$^{13}$C NMR (125 MHz, $(CD_3)_2SO$):** δ = 158.55 (d, *J* = 249.0 Hz), 152.96, 152.40, 150.15, 141.57, 131.18, 130.68 (d, *J* = 9.1 Hz), 129.87 (d, *J* = 3.2 Hz), 129.62, 128.94, 128.80 (d, *J* = 3.1 Hz), 127.84, 127.74, 122.70 (d, *J* = 9.4 Hz), 122.23, 121.12, 118.57 (d, *J* = 25.2 Hz), 108.94 (d, *J* = 141.7 Hz), 14.71. **MS (ESI+):** m/z = 354.14 $[M+H]^+$.

Synthesis of compound **14**; 1-[2-(5-chloro-2-fluorophenyl)-5-methylpyridin-4-yl]-5-methyl-2,3-dihydro-1*H*-1,3-benzodiazol-2-one:

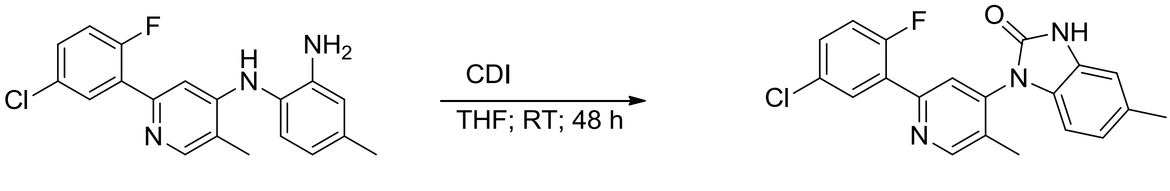

**12** **14**

**12** (300 mg, 0.81 mmol, 1 equiv) was dissolved with CDI (393 mg, 2.42 mmol, 3 equiv) in 30 mL THF. The reaction stirred at RT for 48 h under argon atmosphere. The solvent was evaporated under reduced pressure and the crude product was purified by column chromatography on silica gel (*n*-hexane/EtOAc) to obtain 1-[2-(5-chloro-2-fluorophenyl)-5-methylpyridin-4-yl]-2,3-dihydro-1H-1,3-benzodiazol-2-one as a beige solid. **Yield:** 275 mg (0.75 mmol, 93 %). **$R_f$-Value TLC:** 0.33 (*n*-hexane/EtOAc 1:1). **$^1$H NMR (250 MHz, $(CD_3)_2SO$):** δ = 11.18 (s, 1H), 8.82 (s, 1H), 8.03 (dd, $^3J_{HH}$ = 6.7, $^4J_{HH}$ = 2.8 Hz, 1H), 7.82 (s, 1H), 7.56 (dd, $^3J_{HH}$ = 4.4 Hz, 1H), 7.41 (dd, $^3J_{HH}$ = 8.8 Hz, 1H), 6.93 (s, 1H), 6.83 (d, $^3J_{HH}$ = 8.2 Hz, 1H), 6.71 (d, $^3J_{HH}$ = 8.0 Hz, 1H), 2.33 (s, 3H), 2.21 (s, 3H). **$^{13}$C NMR (125 MHz,

**$(CD_3)_2SO$):** δ = 158.55 (d, $J$ = 249.0 Hz), 152.94, 152.50, 150.06 (d, $J$ = 2.5 Hz), 141.72, 131.52, 131.08, 130.66 (d, $J$ = 9.0 Hz), 129.85 (d, $J$ = 3.3 Hz), 129.05, 128.81 (d, $J$ = 3.0 Hz), 127.78 (d, $J$ = 13.0 Hz), 127.49, 122.58 (d, $J$ = 9.5 Hz), 121.59, 118.56 (d, $J$ = 24.8 Hz), 109.95, 108.13, 20.98, 14.75 ppm. **MS (ESI+):** m/z = 368.14 $[M+H]^+$.

Synthesis of compound **15**; 2-{3-[2-(5-chloro-2-fluorophenyl)-5-methylpyridin-4-yl]-2-oxo-2,3-dihydro-1H-1,3-benzodiazol-1-yl}acetamide:

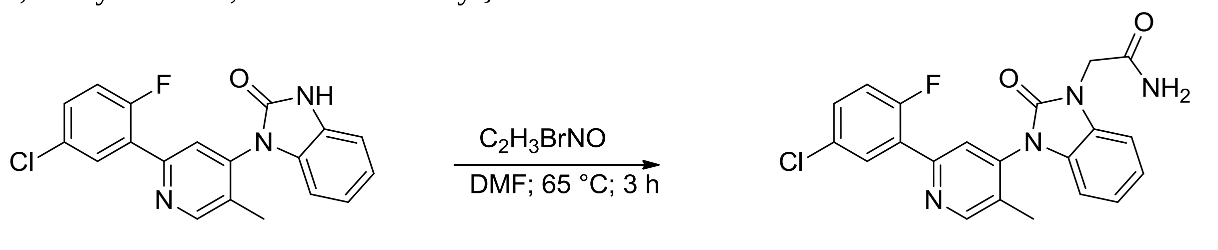

**13** **15**

**13** (245 mg, 0.69 mmol, 1 equiv) was dissolved with 2-bromoacetamide (115 mg, 0.83 mmol, 1.2 equiv) in 40 mL DMF. The reaction stirred at 65 °C for 3h under argon atmosphere. Afterwards the reaction was quenched by the addition of water (200 mL) and the product was extracted with EtOAc. The solvent was removed under reduced pressure and the crude product was further purified by column chromatography on silica gel (n-hexane/EtOAc) to obtain 2-{3-[2-(5-chloro-2-fluorophenyl)-5-methylpyridin-4-yl]-2-oxo-2,3-dihydro-1H-1,3-benzodiazol-1-yl}acetamide as a white solid. **Yield:** 182 mg (0.14 mmol, 63 %). **$R_f$-Value TLC:** 0.21 (*n*-hexane/EtOAc 1:3). **$^1$H NMR (500 MHz, $(CD_3)_2SO$):** δ = 8.86 (s, 1H), 8.05 (dd, $^3J_{HH}$ = 6.8, $^4J_{HH}$ = 2.8 Hz, 1H), 7.87 (d, $^4J_{HH}$ = 1.5 Hz, 1H), 7.71 (s, 1H), 7.57 (ddd, $^3J_{HH}$ = 4.2, $^4J_{HH}$ = 2.8 Hz, 1H), 7.42 (dd, $^3J_{HH}$ = 8.8 Hz, 1H), 7.32 (s, 1H), 7.25–7.19 (m, 1H), 7.17 (dd, $^3J_{HH}$ = 7.6, $^4J_{HH}$ = 1.1 Hz, 1H), 7.07 (td, $^3J_{HH}$ = 7.7, $^4J_{HH}$ = 1.3 Hz, 1H), 6.91 (dd, $^3J_{HH}$ = 7.9, $^4J_{HH}$ = 1.1 Hz, 1H), 4.53 (d, $^4J_{HH}$ = 2.3 Hz, 2 H, $CH_2$), 2.23 (s, 3H). **$^{13}$C NMR (125 MHz, $(CD_3)_2SO$):** δ = 168.43, 158.57 (d, $J$ = 249.4 Hz), 153.10, 151.93, 150.07, 141.46, 131.25, 130.74 (d, $J$ = 9.8 Hz), 130.23, 130.05 , 128.85 (d, $J$ = 1.9 Hz), 128.46, 127.70 (d, $J$ = 12.7 Hz), 122.63 (d, $J$ = 9.6 Hz), 122.24, 121.59, 118.59 (d, $J$ = 24.8 Hz), 108.93, 108.44, 43.28, 14.68 ppm. **MS (ESI+):** m/z = 411.10 $[M+H]^+$. **HPLC:** $t_R$ = 14.171, purity ≥ 95% (UV: 254/280 nm). **HRMS:** m/z calculated for $C_{21}H_{17}ClFN_4O_2^+$: 411.10186 $[M+H]^+$, found: 411.10233 $[M+H]^+$.

Synthesis of compound **16**; 2-{3-[2-(5-chloro-2-fluorophenyl)-5-methylpyridin-4-yl]-6-methyl-2-oxo-2,3-dihydro-1*H*-1,3-benzodiazol-1-yl}acetamide:

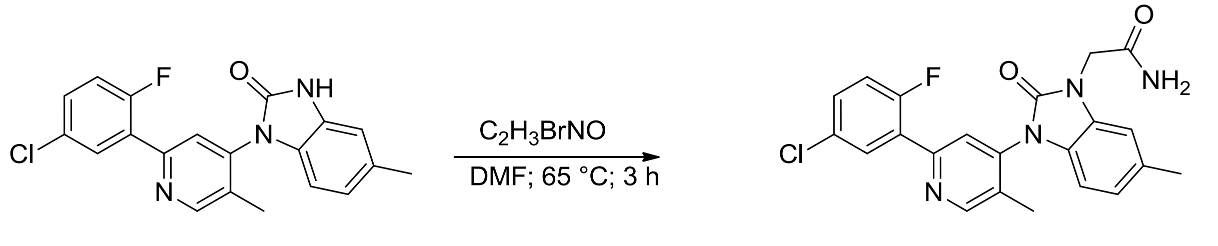

**14** **16**

**14** (275 mg, 0.75 mmol, 1 equiv) was dissolved with 2-bromoacetamide (134 mg, 0.97 mmol, 1.3 equiv) in 27 mL DMF. The reaction stirred at 65 °C for 3h under argon atmosphere. Afterwards the reaction was quenched by the addition of water (200 mL) and the product was extracted with EtOAc. The solvent was removed under reduced pressure and the crude product was further purified by column chromatography on silica gel (n-hexane/EtOAc) to obtain2-{3-[2-(5-chloro-2-fluorophenyl)-5-methylpyridin-4-yl]-6-methyl-2-oxo-2,3-dihydro-1*H*-1,3-benzodiazol-1-yl}acetamide as a white solid. **Yield:** 180 mg (0.424 mmol, 57 %). **$R_f$-Value TLC:** 0.18 (*n*-hexane/EtOAc 1:3). **$^1$H NMR (500 MHz, $(CD_3)_2SO$):** δ = 8.84 (s, 1H), 8.04 (dd,

$^{3}J_{HH}$ = 6.7, $^{4}J_{HH}$ = 2.8 Hz, 1H), 7.84 (d, $^{4}J_{HH}$ = 1.4 Hz, 1H), 7.70 (s, 1H), 7.62–7.50 (m, 1H), 7.42 (dd, $^{3}J_{HH}$ = 8.8 Hz, 1H), 7.31 (s, 1H), 7.03 (d, $^{4}J_{HH}$ = 1.5 Hz, 1H), 6.95–6.85 (m, 1H), 6.79 (d, $^{3}J_{HH}$ = 8.0 Hz, 1H), 4.49 (d, $^{4}J_{HH}$ = 3.0 Hz, 2H), 2.35 (s, 3H), 2.22 (s, 3H). **$^{13}$C NMR (125 MHz, $(CD_3)_2SO$:** δ = 168.91, 159.03 (d, *J* = 249.0 Hz), 153.54, 152.49, 150.45 (d, *J* = 2.6 Hz), 142.06, 132.07, 131.61, 131.17 (d, *J* = 9.1 Hz), 130.78, 130.32 (d, *J* = 3.8 Hz), 129.31 (d, *J* = 3.0 Hz), 128.15 (d, *J* = 13.0 Hz), 126.80, 122.98 (dd, *J* = 9.8, 4.6 Hz), 122.48, 119.05 (d, *J* = 24.9 Hz), 109.81, 108.68, 43.67, 21.52, 15.18. **MS (ESI+):** m/z = 447.09 [M+ Na] $^{+}$. **HPLC:** $t_R$ = 14.875, purity ≥ 95% (UV: 254/280 nm). **HRMS:** m/z calculated for $C_{22}H_{19}ClFN_4O_2^{+}$: 425.11751 $[M+H]^{+}$, found: 425.11825 $[M+H]^{+}$.

<sup></sup>

[1]H NMR of Compound **15** (THAI11).

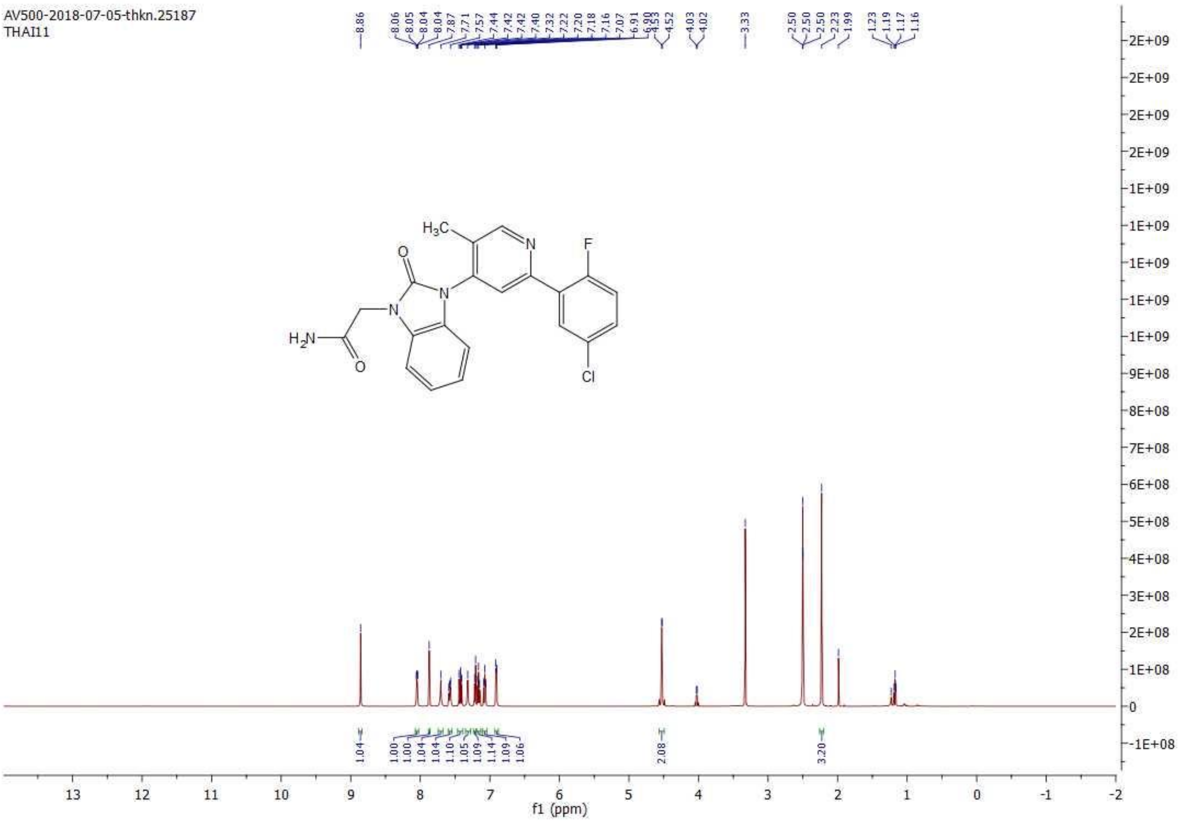

[13]C NMR of Compound **15** (THAI11).

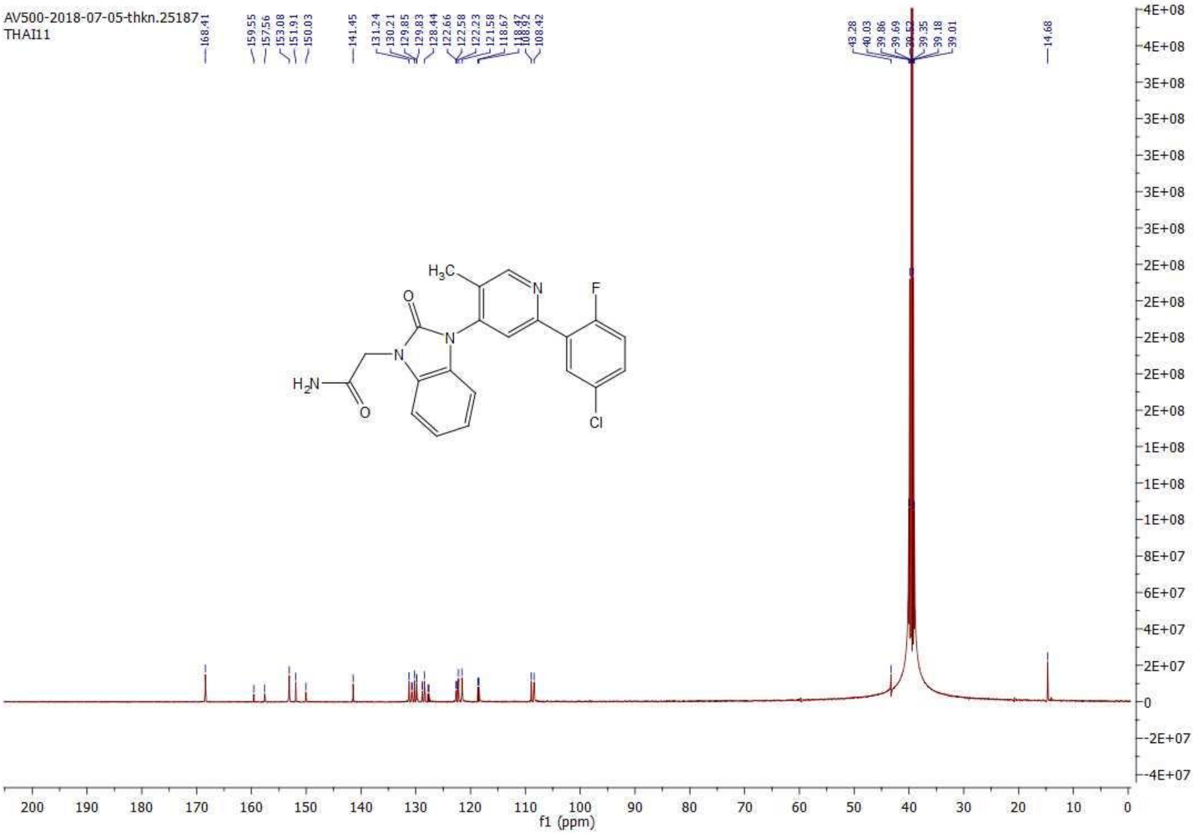

[1]H NMR of Compound **16** (THAI14).

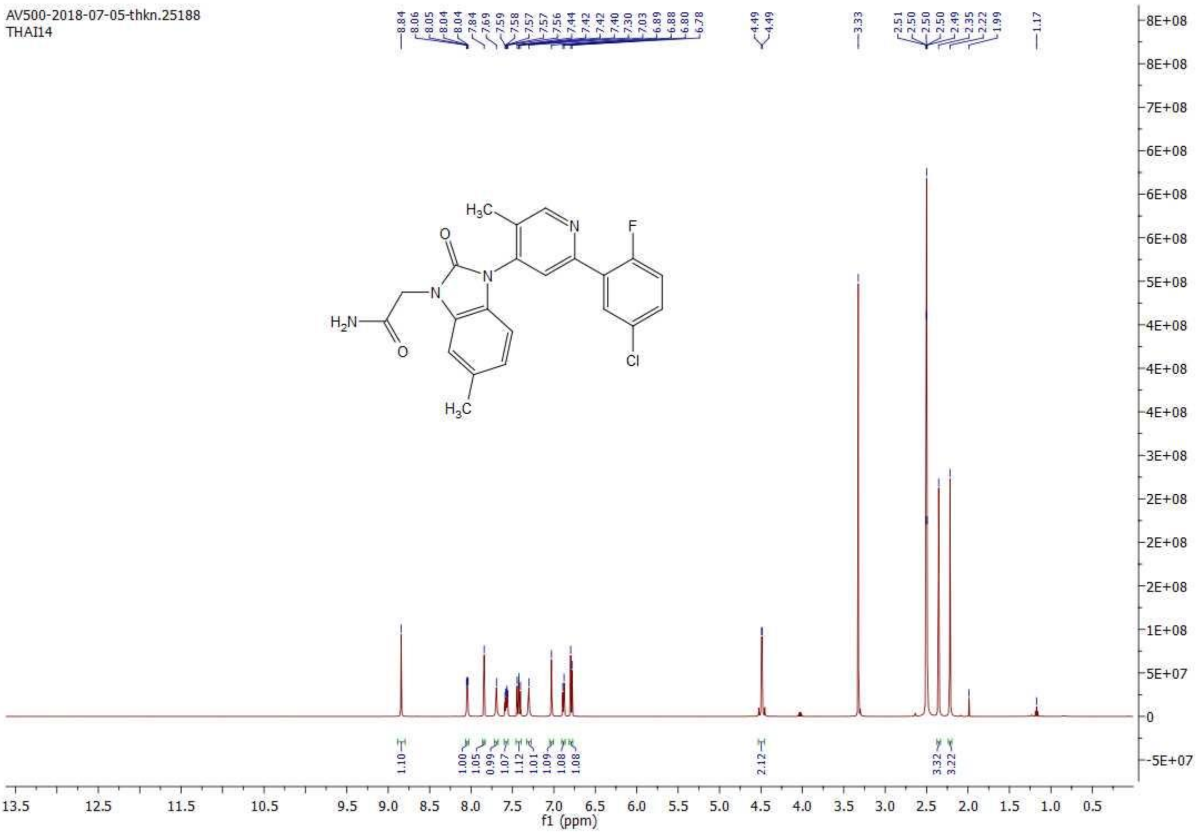

[13]C NMR of Compound **16** (THAI14).

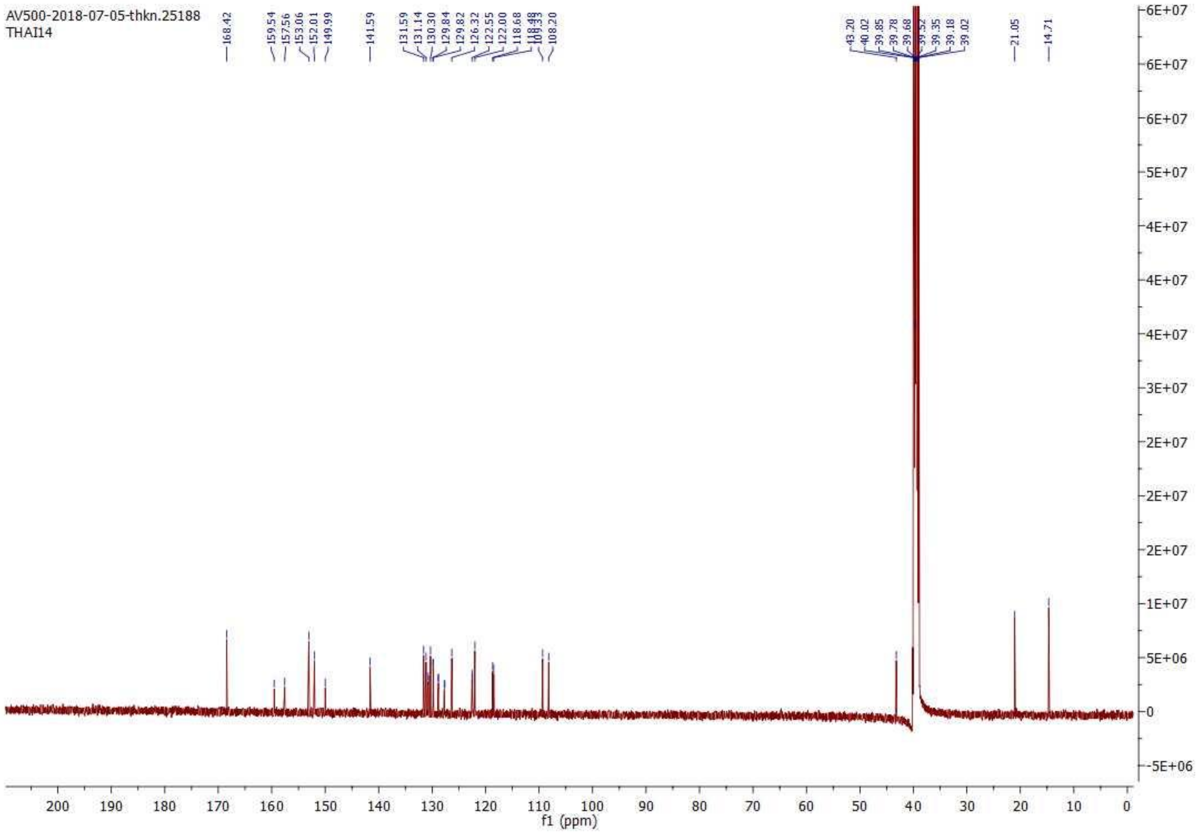
